# MitoSAM-dependent lipoylation controls postnatal heart development via metabolic remodeling

**DOI:** 10.64898/2026.04.20.719537

**Authors:** Anastasia Rumyantseva, Alissa Wilhalm, William Carter, Trevor S. Tippetts, Marco Moedas, Florian A. Rosenberger, David Moore, Ákos Végvári, Yvonne Hinze, Linden Muellner-Wong, David Alsina, Rolf Wibom, Rainah Winston, Thomas P. Mathews, Lars H. Lund, Daniel C. Andersson, Gianluigi Pironti, Anna Wedell, Ralph J. DeBerardinis, Christoph Freyer, Anna Wredenberg

**Author notes:** Senior author (Christoph Freyer, Anna Wredenberg).

## Abstract

The neonatal heart undergoes a rapid metabolic transition from fetal glycolysis to oxidative phosphorylation, requiring coordinated metabolic remodeling. Mechanisms driving this transition remain unclear. Here, we demonstrate that sufficient mitochondrial S-adenosylmethionine (mitoSAM), imported via the solute carrier *Slc25a26*, is essential for this shift by sustaining the lipoylation of 2-oxoacid dehydrogenases, critical for TCA cycle activation. Proteomic and metabolomic profiling revealed that reduced mitoSAM availability impaired lipoylation, blocking TCA cycle function and restricting nucleotide synthesis, while mitochondrial gene expression and respiratory capacity remained largely intact. In vivo EdU labeling showed persistent cardiomyocyte proliferation imposing further strain on nucleotide pools. Supplementation with medium-chain triglycerides during the suckling-to-weaning transition restored metabolic function and normalized cardiac growth and morphology. Our data reveal a critical developmental window in which mitoSAM-dependent lipoylation ensures heart maturation.

---

During the early postnatal period the mammalian heart undergoes profound metabolic remodeling to meet the energetic demands of continuous contractile function. This transition is shaped by rising oxygen availability at birth and successive shifts in nutrient supply: from placental glucose and lactate in utero, to lipid-rich milk, and later carbohydrate-based solid food. Simultaneously, a switch towards oxidative metabolism occurs, placing mitochondria at the center of cardiomyocyte maturation, as the adult myocardium relies primarily on fatty acid β-oxidation for ATP production (*1–3*). Yet how these transcriptional and metabolic programs are coordinated, and which metabolites are critical for this developmental phase, remain poorly understood.

S-adenosylmethionine (SAM) is a universal biological methyl group donor synthesized from methionine and ATP, driving a wide range of methylation and radical-based reactions across cellular compartments (*4*). Within mitochondria, SAM (mitoSAM) supports two distinct processes: (i) classical methylation reactions, essential for mitochondrial gene expression and coenzyme Q (CoQ) biosynthesis, and (ii) radical-mediated synthesis of lipoic acid (LA) by lipoic acid synthetase (LIAS) (*5*). LA is a rare, highly conserved posttranslational lysine modification within mitochondria, required for the glycine cleavage system H protein (GCSH) and the four oxoacid dehydrogenase complexes: pyruvate dehydrogenase (PDHc), branched-chain α-ketoacid dehydrogenase (BCKDHc), 2-oxoadipate dehydrogenase (OADHc), and α-ketoglutarate dehydrogenase, also known as 2-oxoglutarate dehydrogenase (OGDHc) (*6*). Covalently bound to the E2 subunits, LA acts as a flexible ‘swinging arm’ that channels reaction intermediates between active sites, enabling coordinated redox chemistry and acyl-group transfer (*7–10*).

As a substrate for mitochondrial methyltransferases and radical SAM enzymes, mitoSAM coordinates gene expression, oxidative phosphorylation, and TCA cycle activity within mitochondria. Yet how mitoSAM availability shapes these core metabolic pathways in vivo remains underexplored. This question is particularly pertinent in the heart, as it is acutely sensitive to primary mitochondrial dysfunction (*11–13*). Moreover, the neonatal heart passes through a window of developmental vulnerability, during which nutrient supply and cofactor availability can have disproportionate and lasting effects on organ function (*14–19*). Notably, metabolic remodeling also occurs in the adult heart during pathological states such as heart failure, where altered mitochondrial substrate utilization and TCA cycle activity are increasingly recognized as contributing factors (*20*, *21*). Here, we probe how mitoSAM availability constrains metabolic maturation of the postnatal heart, by targeting mitoSAM availability in heart- and skeletal muscle-specific knockout mouse model.

## Slc25a26 is enriched in the cardiomyocyte lineage

The solute carrier 25 (SLC25) family comprises over 50 mitochondrial transporters that shuttle a wide range of metabolites across the inner mitochondrial membrane; many of which are considered to be ubiquitously expressed (*22–24*). Among these, *SLC25A26* encodes the only known SAM carrier, which imports cytosolic SAM into the mitochondrial matrix (Fig. 1A) (*5*, *25–27*). Analysis of a publicly available single-cell RNA sequencing dataset covering 20 mouse tissues (*28*) revealed striking tissue-specific heterogeneity, and *Slc25a26* expression was particularly high in cardiomyocytes across the entire dataset (Fig. 1B, fig. S1A). To account for differences in mitochondrial content across tissues, we compared *Slc25a26* expression in the cardiomyocyte lineage to other mitochondrial SLC25 carriers. Only *Slc25a3* (phosphate transporter) and *Slc25a4* (ADT/ATP translocase) reached comparable expression levels in cardiomyocytes, suggesting an unusually high demand for mitoSAM in this lineage (Fig. 1C, fig. S1B).

**Fig. 1.**
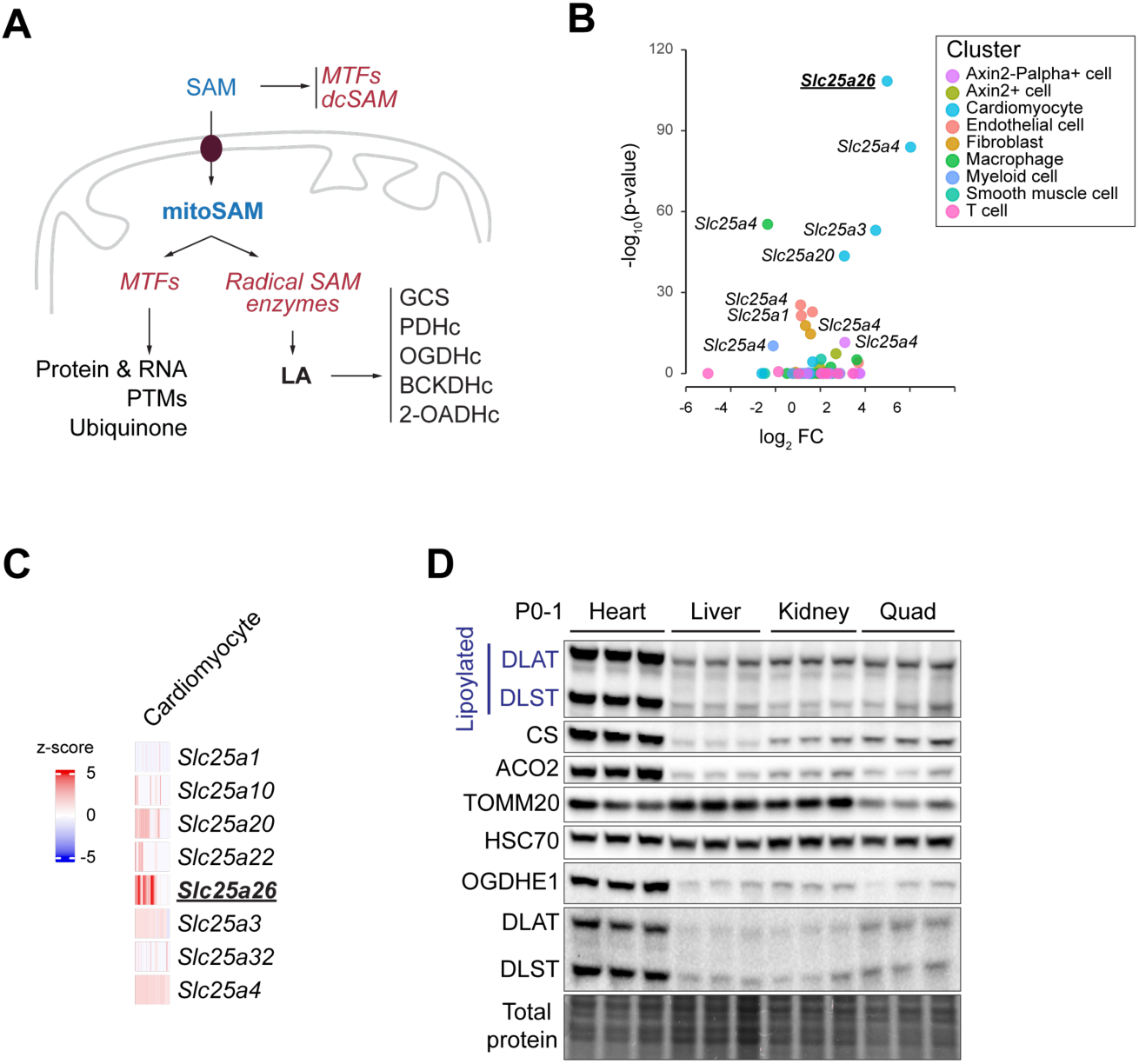
Slc25a26 shows specific enrichment in cardiomyocyte lineage. (**A**) Schematic diagram of SAM functions and its import into mitochondria. SAM is synthesized in the cytoplasm and used either as methyl group donor by MTFs, decarboxylated (dcSAM), or as substrate for radical SAM enzymes for the synthesis of i.e. lipoic acid; an obligatory co-factor of the glycine cleavage system H protein (GCS), and the pyruvate dehydrogenase (PDHc), α-ketoglutarate dehydrogenase (OGDHc), branched-chain α-ketoacid dehydrogenase (BCKDHc), and 2-oxoadipate dehydrogenase (2-OADHc) complexes. (**B**) Volcano plot depicting the log_2_FC in differential expression of selected SLC25 family members in various clusters compared to the total population. Data were calculated from using a publicly available data set using a Wilcox’s test with Bonferroni correction. (**C**) Heat map showing indicated *Slc25a* family members expression in cardiomyocyte lineage. (**D**) Immunoblot of control (Ctrl) tissue lysates in heart, liver, kidney, and quadriceps (Quad) at P0-1(n=3) with indicated antibodies. HSC70 was used as the loading control. SAM, s-adenosylmethionine; MTFs, methyltransferases; PTMs, posttranslational modifications; LA, lipoic acid.

The enrichment of *Slc25a26* in cardiomyocytes likely reflects their adaptation to the postnatal shift toward oxidative metabolism. We therefore asked whether mitoSAM levels change during this transition. Although cellular SAM pools have been shown to fluctuate markedly during early postnatal development (*29–31*), targeted metabolomics revealed that mitoSAM levels remained remarkably unaltered from birth to weaning (fig. S1C). This stability suggests tight homeostatic control during the rapid expansion of mitochondrial mass and OXPHOS activity during this period (*3*, *32*).

Given the central role of the tricarboxylic acid (TCA) cycle in energy metabolism, we next examined its postnatal regulation. TCA cycle protein abundance increased during the neonatal-to-weaning period (fig. S1D). We then assessed mitoSAM-dependent lipoylation, required for the activity of PDHc and OGDHc. Both lipoylation of their E2 subunits (DLAT and DLST, respectively) and apo-forms were markedly higher in perinatal hearts than in other tissues (Fig. 1D) and increased further between postnatal day P4 and P14 (fig. S1D), coinciding with the postnatal increase in oxidative metabolism (*3*, *32*). Together, these findings identify *Slc25a26* as highly enriched in the cardiomyocyte lineage, reveal that mitoSAM levels are stably maintained despite postnatal remodeling, and show coordinated upregulation of TCA cycle protein abundance and lipoylation during postnatal heart maturation.

## Loss of Slc25a26 impairs cardiac lipoylation and accumulates α-ketoglutarate before mitochondrial gene expression is affected

The high expression of *Slc25a26* in cardiomyocytes, along with the postnatal increase in lipoylation suggest an elevated demand for mitoSAM during heart maturation. Targeting *Slc25a26* therefore allows us to dissect the specific contributions of mitoSAM to methylation and LA biosynthesis, and their subsequent roles in shaping cardiac metabolism and maturation.

Biallelic variants in *SLC25A26* in pediatric and adult patients are associated with lactic acidosis, episodes of acute cardiopulmonary failure, progressive muscle weakness, and, in severe cases, neonatal mortality (*33–36*). In model organisms ranging from yeast to mice, complete loss of *Slc25a26* is incompatible with life and causes profound metabolic alterations, underscoring a conserved requirement for mitoSAM (*27*, *36–38*).

To directly test the role of mitoSAM transport in the postnatal heart, we crossed *Slc25a26^LoxP/LoxP^* mice (*27*) with transgenic animals expressing *Cre* under the *Ckmm*-promoter from day E13.5 (*39*). This resulted in efficient deletion in heart and skeletal muscle but not liver (Fig. 2A, fig. S2A). These conditional KO mice (hereafter referred to as *Slc25a26^KO^*) were born with no overt phenotype and were indistinguishable from control littermates at P7, with no evidence of cardiomyopathy (fig. S2B to D).

**Fig. 2.**
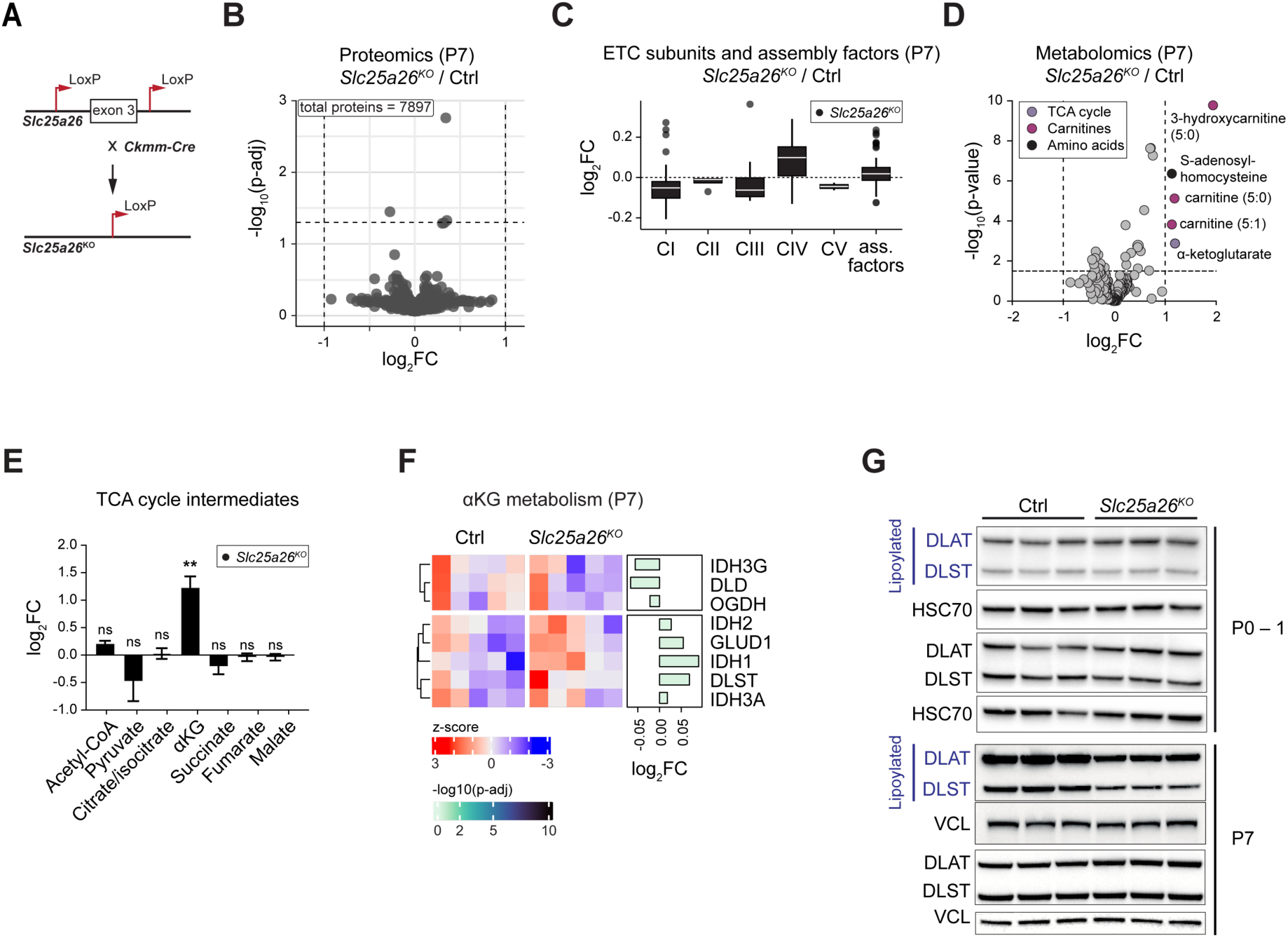
Lipoylation defect precedes mitochondrial gene expression impairment in *Slc25a26^KO^*model. (**A**) Schematic diagram of *Slc25a26^KO^* mice breeding. (**B**) LC-MS/MS label-free whole heart proteomics, volcano plot of *Slc25a26^KO^* compared with Ctrl at P7 (n=5 per genotype). (**C**) Boxplots of ETC subunits and assembly factors in *Slc25a26^KO^*heart proteome at P7, log_2_FC, relative to Ctrl. (**D**) Volcano plot of untargeted metabolome data, showing differentially abundant metabolites in *Slc25a26^KO^* versus Ctrl hearts at P7 (n=6 per genotype). Metabolites are colored according to categories. (**E**) Log_2_FC of TCA cycle intermediates from untargeted metabolomics in *Slc25a26^KO^* hearts at P7 relative to Ctrl. (**F**) Heat map with relative z-score of heart proteomics data, showing proteins in αKG metabolism. (**G**) Immunoblot of Ctrl and *Slc25a26^KO^* heart lysates, depicting lipoylated and apo-forms of DLAT and DLST at P0-1 and P7, vinculin (VCL) or HSC70 were used as the loading controls (n=3 per genotype). Data in (E) presented as log_2_FC ± s.e.m., two-tailed unpaired Student’s *t*-test; ns = not significant, **p <0.01.

We recently demonstrated that mitoSAM-dependent methylation promotes mitochondrial ribosome assembly and translation in mouse embryonic fibroblasts (MEFs) and skeletal muscle (*40*). Surprisingly, P7 hearts showed no major changes in electron transport chain (ETC) subunit levels (Fig. 2B and C, fig. S2E, data S1), aside from a mild increase in *mtNd3* transcripts (fig. S2F). Mitochondrial stress markers *Atf4*, *Atf5*, and *Chop* were not induced (fig. S2G), and label-free LC-MS/MS proteomics revealed no significant alterations in ETC accessory components, stress proteins, or the cardiac contractile proteome (fig. S3A and B). These findings indicate that loss of mitoSAM transport does not impair mitochondrial gene expression or trigger stress signaling in the early postnatal heart.

Given the absence of apparent cardiomyopathy at P7, we next asked whether early metabolic changes could reveal vulnerabilities that precede structural or transcriptional defects. Untargeted metabolomics of P7 *Slc25a26^KO^* and control hearts identified only five significantly altered metabolites (Fig. 2D, fig. S3C, data S2). In addition to several C5-carnitines, reported to be byproducts of incomplete branched-chain amino acid oxidation (*41*), we observed a striking two-fold increase in α-ketoglutarate (αKG) in *Slc25a26^KO^* hearts. Levels of other TCA cycle intermediates and 2-hydroxyglutarate (2-HG) were unchanged, as were protein abundances of isocitrate dehydrogenase, glutamate dehydrogenase, and OGDHc subunits (Fig. 2E and F), pointing to a possible defect in OGDHc activity.

OGDHc function depends on lipoylation of its E2 subunit (DLST), and we therefore examined protein lipoylation during early postnatal development. At birth, *Slc25a26^KO^* hearts displayed normal lipoylation, but levels began to decline at P4 and fell progressively throughout the first postnatal week (Fig. 2G, fig. S3D), coinciding with the developmental shift to oxidative metabolism (*3*, *32*). This occurred despite only a mild reduction in mitoSAM levels at P7 (fig. S3E), suggesting that increased metabolic demand during maturation unmasks the sensitivity of the lipoylation pathway to reduced mitoSAM supply. Notably, mitochondrial s-adenosylhomocysteine (mitoSAH) levels were elevated, consistent with increased SAH levels in the untargeted metabolomic data (Fig. 2D, fig. S3E).

Thus, cardiomyocyte-specific loss of *Slc25a26* triggers a progressive defect in protein lipoylation that arises before detectable changes in mitochondrial gene expression or ETC subunits abundance. These data identify lipoylation as an early and uniquely mitoSAM-sensitive node in cardiac metabolism during postnatal maturation.

## Slc25a26 ablation results in cardiomyopathy

The early and progressive loss of lipoylation in *Slc25a26^KO^* hearts prompted us to examine the physiological consequences during postnatal maturation. Although *Slc25a26^KO^* mice appeared healthy at birth, they were prone to sudden death shortly after P14, with a median lifespan of 15 days (Fig. 3A). By P14, heart-weight-to-body-weight (HW/BW) ratio was significantly elevated, reflecting increased cardiac mass (fig. S4A to C). Histological analysis revealed vacuolar degeneration of cardiomyocytes (Fig. 3B). Moreover, the cardiomyocyte cell size was significantly smaller in *Slc25a26^KO^* hearts in wheat germ agglutin-stained sections (Fig. 3B and C).

**Fig. 3.**
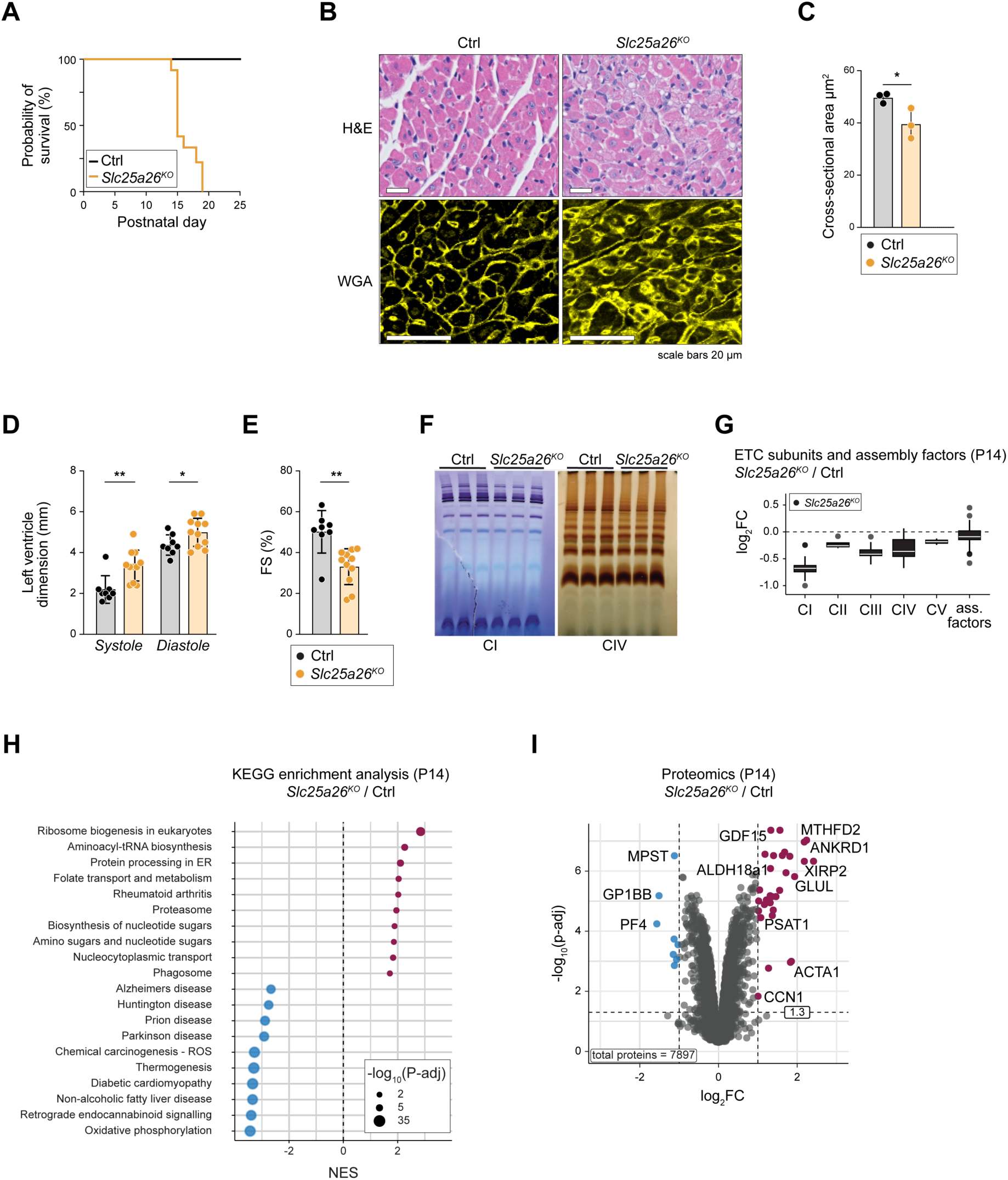
Slc25a26 loss results in cardiomyopathy. (**A**) Kaplan-Meier survival curve of *Slc25a26^KO^*mice (n=11 mice). (**B**) Representative images of H&E and WGA staining of heart sections of Ctrl and *Slc25a26^KO^* mice at P14 (n=3 per genotype, scale bars =20 μm). (**C**) Quantification of cross-sectional area of cardiomyocytes in Ctrl and *Slc25a26^KO^* mice (n=3 per genotype). (**D**) Echocardiographic analysis of Ctrl (n=8) and *Slc25a26^KO^* mice (n=11) at P12, indices of the LV dimensions in diastole and systole. (**E**) Fractional shortening of LV in Ctrl (n=8) and *Slc25a26^KO^* (n=11) hearts at P12. (**F**) Enzymatic in-gel activity staining for CI and CIV from digitonin-solubilized cardiac mitochondria at P14, separated by BN-PAGE (n=3 per genotype). (**G**) LC-MS/MS label-free whole heart proteomics. Boxplots of ETC subunits and assembly factors in *Slc25a26^KO^* hearts relative to controls at P14. (n=5 for Ctrl, n=4 for *Slc25a26^KO^*). (**H**) KEGG pathway enrichment analysis of significantly up- (purple) or downregulated (blue) proteins in *Slc25a26^KO^* hearts. (**I**) Volcano plot of differentially expressed proteins in *Slc25a26^KO^* hearts compared with Ctrl at P14, significantly changed proteins (log_2_FC > 1 and log_2_FC < – 1, p-adj < 0.05) are shown in purple (increased) and blue (decreased). Data in (C, D and E) are mean ± SD, each dot represents individual mouse, two-tailed unpaired Student’s *t*-test, **p <0.01, *p < 0.05.

Echocardiography was performed at P12 to avoid anesthesia complications. *Slc25a26^KO^* hearts displayed enlarged left ventricular dimensions, reduced fractional shortening, and thinning of the interventricular septum, while posterior wall thickness remained unchanged (Fig. 3D and E, fig. S4D). These features indicate the early-onset cardiomyopathy as a direct consequence of mitoSAM deficiency.

Given the severity of the phenotype, we anticipated profound respiratory chain defects. However, in-gel activity assays of complexes I and IV revealed that both supercomplex formation and activity were only mildly affected in *Slc25a26^KO^* hearts (Fig. 3F). Spectrophotometric measurements confirmed moderate decreases in CI/CIII- and CII/CIII-linked activities (∼60% of control), likely reflecting reduced CoQ levels (fig. S4E and F), while CIV activity was decreased by only ∼15% (fig. S4E).

Consistent with these findings, steady-state levels of mitochondrial transcripts remained mostly unchanged (fig. S4G), and proteomics detected only a modest reduction in complex I subunits (log₂FC ≈ – 0.7), with other ETC complexes largely unaffected (Fig. 3G, fig. S4H, data S1). KEGG pathway analysis of upregulated proteins highlighted ribosome biogenesis, folate transport, and aminoacyl-tRNA biosynthesis (Fig. 3H). At the individual protein level, this was reflected by induction of mitochondrial and cardiac stress markers (GDF15, ANKRD1, MTHFD2) together with enzymes of amino acid and one-carbon metabolism (MTHFD2, PSAT1, GLUL, ALDH18A1), pointing towards activation of stress-response pathways (*42*) (Fig. 3I).

Thus, despite retaining near-normal mitochondrial gene expression and only modest reductions in ETC capacity *Slc25a26^KO^* hearts present with a rapidly progressive cardiomyopathy and early lethality. Taken together, this points towards alternative metabolic defects as a driver of pathology.

## Slc25a26 ablation remodels heart metabolism and disrupts mitochondrial enzyme function

Early metabolic changes at P7, together with moderate ETC impairment at P14, prompted us to ask whether broader metabolic pathways might be disrupted in *Slc25a26^KO^* hearts. Untargeted metabolomics of P14 hearts revealed profound changes, with 51 of 221 detected metabolites significantly altered in *Slc25a26^KO^* hearts (Fig. 4A, fig. S5A and B, data S2). The accumulation of αKG and branched-chain ketoacids together with reduced succinate, aspartate, and nucleotide pools were consistent with impaired activity of the lipoylation-dependent 2-oxoacid dehydrogenases OGDHc, BCKDHc, and OADHc (Fig. 4A and B). Additional alterations across carnitines, fatty acids, and phospholipid intermediates likely reflected secondary consequences of this block (fig. S5B). Indeed, lipoylated DLAT and DLST were nearly absent in *Slc25a26^KO^* hearts, while lipoylation was preserved in quadriceps from the same mice (Fig. 4C, fig. S5C), confirming a heart-specific defect in mitochondrial enzyme function.

**Fig. 4.**
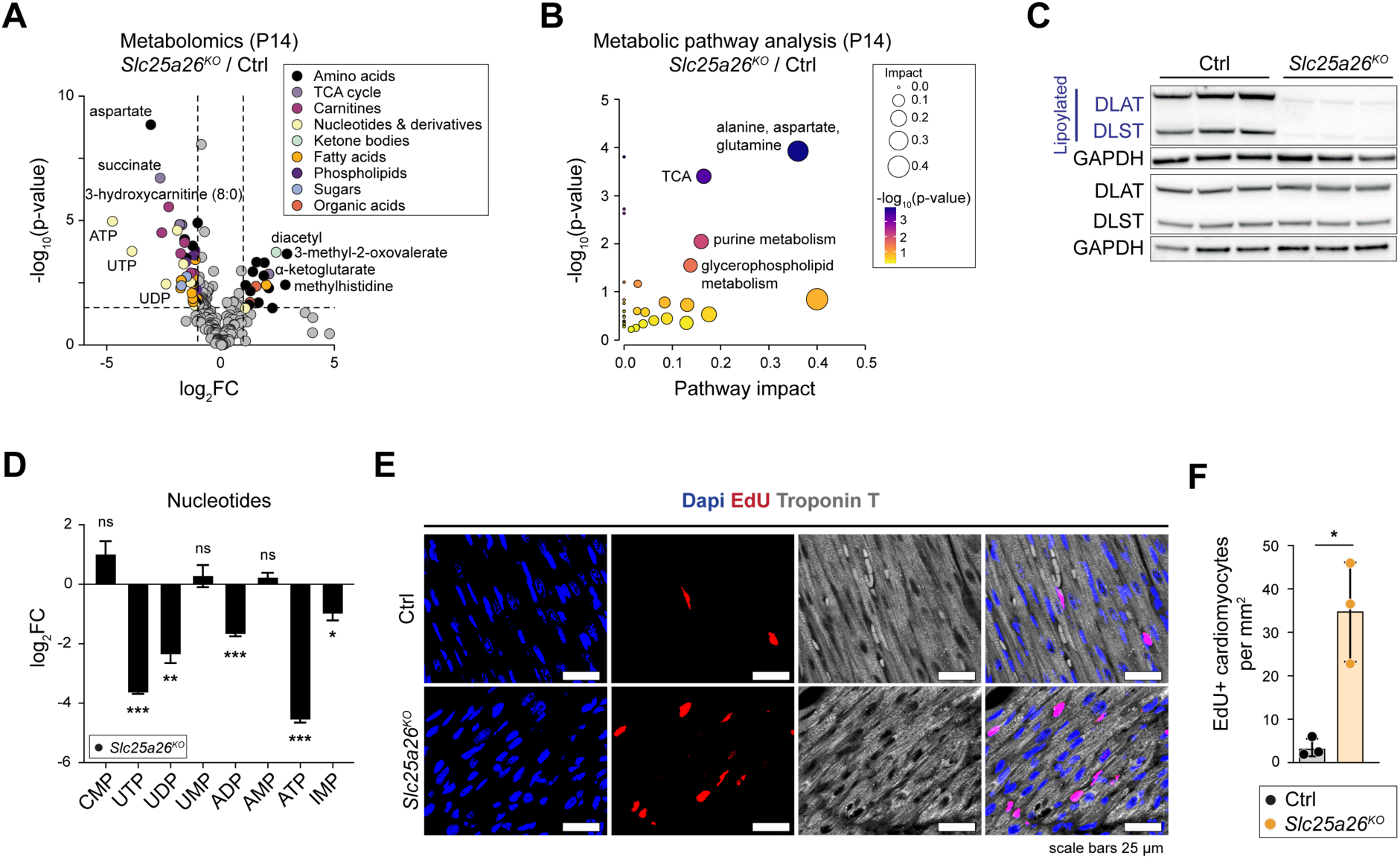
Slc25a26 loss triggers metabolic perturbation and prevents cell-cycle exit. (**A**) Volcano plot of whole heart untargeted metabolome, showing differentially abundant metabolites of *Slc25a26^KO^*comparing to Ctrl at P14. Significantly altered metabolites (log_2_FC > 1 and log_2_FC < –1, p-value < 0.05) are colored according to categories (n=7 for Ctrl, n=5 for *Slc25a26^KO^*). (**B**) Metabolic pathway analysis of differentially altered metabolites in *Slc25a26^KO^* mice compared with Ctrl. (**C**) Immunoblot of Ctrl and *Slc25a26^KO^* heart lysates, depicting lipoylated and apo-forms of DLAT and DLST at P14, GAPDH used as the loading control (n=3 per genotype). (**D**) Log_2_FC of nucleotides detected in the untargeted metabolome of *Slc25a26^KO^* hearts compared with Ctrl at P14. (**E**) Representative immunofluorescent images of P14 heart sections as indicated, depicting EdU incorporation (EdU), cardiomyocytes (anti-cardiac troponin T) and nuclei (DAPI). (**F**) Quantification of (E) EdU^+ve^ cardiomyocytes. Data in (D) presented as log_2_FC ± s.e.m., two-tailed unpaired Student’s *t*-test. Data in (E) are mean ± SD, each dot represents an individual mouse, two-tailed unpaired Student’s *t*-test, ns = not significant, ***p <0.001, **p <0.01, *p < 0.05.

## Loss of lipoylation alters cysteine oxidation of 2-oxoacid dehydrogenases

Several notable antioxidants were decreased in *Slc25a26^KO^*mice, suggesting impaired reductive-oxidation homeostasis (fig. S5D). We therefore performed redox proteomics to assess the oxidation state of cysteine residues (*43*). Only nine cysteine sites on mitochondrial proteins showed increased oxidation in *Slc25a26^KO^* hearts (fig. S5E, data S3). Interestingly, we detected all known lipoate-modified peptides, except for GCSH. Notably, these sites were located near the conserved lipoylated lysines of OGDHc and PDHc subunits (fig. S5F), suggesting that lipoate stabilizes local protein structure, shielding adjacent cysteines from oxidation. Similar protective effects have been proposed in *ex vivo* studies (*44*, *45*). Additionally, we observed opposing redox shifts in malate dehydrogenase 2 (MDH2, oxidized) and glutamate/oxaloacetate transaminase 2 (GOT2, reduced), two core malate/aspartate shuttle enzymes essential for cardiac aspartate synthesis (*46*). However, the absence of widespread oxidation suggests that oxidative stress is not the primary driver of cardiac failure in these mice.

## Slc25a26 deficiency triggers metabolic perturbation/alteration and immature cardiac identity

Given the central role of lipoylated dehydrogenases in TCA cycle flux, we next examined the downstream metabolic consequences. Nucleotide profiling revealed a pronounced depletion of ATP, UTP, and UDP, alongside reduced aspartate, an essential substrate for nucleotide biosynthesis, and purine intermediates such as IMP and succinyladenosine (Fig. 4D, fig. S6A and B). Although adenine pools are often reduced in heart failure (*47*, *48*), *Slc25a26^KO^* hearts showed decreases not only in precursors but also in degradation products such as adenosine and xanthine (fig. S6A and B). Proteomics further revealed profound changes in nucleotide metabolism, including upregulation of enzymes in the de novo purine and pyrimidine biosynthetic pathways (IMPDH1, IMPDH2, TYMS, CTPS1) and increased synthesis of nucleotide sugars (Fig. 3H, fig. S6C and D), suggesting a compensatory response.

De novo nucleotide biosynthesis is energetically demanding, and cardiomyocytes cease proliferation after P7 (*3*) relying on the nucleotide salvage pathway (*49*). Thus, we hypothesized that this metabolic profile reflected persistent proliferation. Consistent with this, in vivo EdU incorporation confirmed that P14 control hearts had largely exited the cell cycle, whereas *Slc25a26^KO^* hearts retained a significantly higher fraction of proliferating cardiomyocytes (Fig. 4E and F). In addition, transcript levels of mature sarcomeric gene isoforms and pivotal calcium-handling genes (*Atp2a2*, *Ryr2*, *Kcnip2*) were reduced, while fetal isoforms and heart failure markers (*Nppa*, *Nppb*) were elevated (fig. S7A and B). Proteomics confirmed this shift, showing downregulation in the cardiac muscle contraction pathway (fig. S7C).

αKG has been reported to epigenetically stimulate cardiomyocyte proliferation in neonates and post-myocardial infarction via αKG-dependent dioxygenases (*16*, *50*). Although αKG was increased four-fold in P14 *Slc25a26^KO^* hearts (fig. S7D), this was matched by elevated 2-HG (fig. S7D and E), a known inhibitor of αKG-dependent dioxygenases (*51*). In line with this, global 5hmC levels remained unchanged at P7 and P14 (fig.S7F).

*Slc25a26* ablation therefore drives a metabolome-first collapse through the loss of lipoylation disrupting the TCA cycle, prolonging the proliferative window and ultimately depleting the nucleotide pool. Although localized epigenetic effects cannot be excluded, the unchanged global 5hmC profile suggests that the dual bioenergetic and biosynthetic burden stems primarily from metabolic dysfunction.

## Lipoylation-dependent TCA flux is the primary bottleneck in Slc25a26^KO^ hearts

Mice undergo a critical suckling-weaning transition around P12, when milk rich in medium-chain triglycerides (MCTs) is steadily replaced by a carbohydrate-based chow diet (*52*). This coincides with rising cardiac energy demands, creating a dual energetic challenge of: (i) declining MCT supply, and (ii) increased reliance on pyruvate and glutamate oxidation triggered by metabolic remodeling (*20*, *53*). *Slc25a26^KO^*hearts, however, are likely unable to meet this shift due to blockage at the lipoylation-dependent complexes (fig. S7G).

To define this bottleneck, we measured substrate-specific ATP production in isolated mitochondria. Rates from succinate and acylcarnitines (medium- or long-chain) were preserved or only mildly affected but severely impaired for substrates requiring the lipoylated enzymes PDHc (pyruvate, malate) or OGDHc (αKG, glutamate) (Fig. 5A). Thus, while OXPHOS itself remained functional, loss of flux through lipoylation-dependent dehydrogenases rendered *Slc25a26^KO^* hearts unable to adapt to changing carbon sources when metabolic flexibility is critical.

**Fig. 5.**
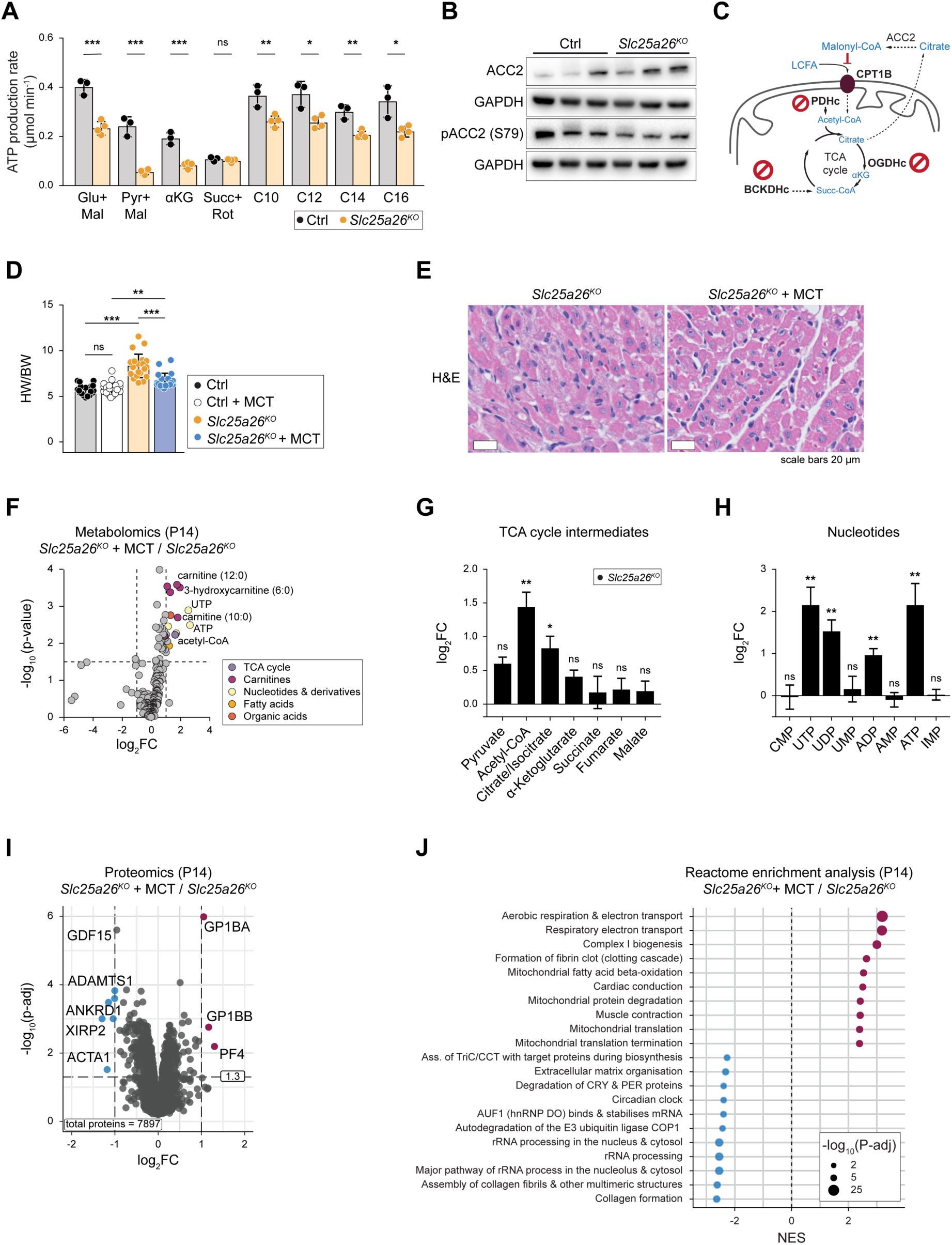
Medium-chain triglyceride-enriched diet mitigates metabolic collapse in Slc25a26^KO^. (**A**) *Ex vivo* ATP production assay in Ctrl (n=3) and *Slc25a26^KO^* (n=4) P14 mitochondria supplied with indicated substrates. (**B**) Immunoblot of Ctrl and *Slc25a26^KO^* heart lysates, depicting total and phosphorylated (S79) ACC. GAPDH was used as a loading control. (**C**) Scheme of impaired cardiac substrate utilization in *Slc25a26^KO^* mice at P14. (**D**) Heart weight (HW) normalized to body weight (BW) of P14 mice with MCT diet (n=21, 26, 20, 18 for Ctrl, Ctrl+MCT, *Slc25a26^KO^* and *Slc25a26^KO^* +MCT, respectively). (**E**) H&E staining of heart sections of *Slc25a26^KO^*and *Slc25a26^KO^* +MCT mice at P14. (**F**) Volcano plot of untargeted metabolome data, showing differentially abundant metabolites in hearts of P14 *Slc25a26^KO^* mice fed on MCT-enriched versus normal chow. Significantly altered metabolites (log_2_FC > 1 and log_2_FC < –1, p-value < 0.05) are colored according to categories (n=5 for *Slc25a26^KO^*, n=6 for *Slc25a26^KO^* +MCT). (**G**) Log_2_FC of TCA cycle intermediates detected in the untargeted heart metabolome of P14 *Slc25a26^KO^* mice fed on MCT-enriched versus normal chow. (**H**) Log_2_FC of nucleotides detected in the untargeted heart metabolome of P14 *Slc25a26^KO^* mice fed on MCT-enriched versus normal chow. (**I**) LC-MS/MS label-free whole heart proteomic analysis. Volcano plot of differentially expressed proteins in hearts from P14 *Slc25a26^KO^*mice fed on MCT-enriched versus normal chow. Significantly changed proteins (log_2_FC > 1 and log_2_FC < – 1, p-adj < 0.05) are shown in purple (increased) and blue (decreased). (**J**) Reactome pathway enrichment analysis of significantly up- (purple) or downregulated (blue) proteins in *Slc25a26^KO^* +MCT hearts. Data in (A and D) are presented as mean ± SD; each dot represents individual mouse. Two-tailed unpaired Student’s *t*-test (A) and two-way ANOVA with Tukey’s multiple comparisons test (D). Data in (G and H) presented as log_2_FC ± s.e.m., statistical significance was determined using two-tailed unpaired Student’s *t*-test; ns = not significant, *p<0.05; **p<0.01; ***p<0.001.

Interestingly, *Slc25a26^KO^* hearts also displayed decreased levels of phosphorylated acetyl-CoA carboxylase (pACC), suggestive of increased malonyl-CoA production (Fig. 5B and C). Malonyl-CoA is a potent inhibitor of CPT1B, the rate-limiting transporter for long-chain fatty acid oxidation (*54*, *55*), potentially restricting the utilization of long-chain fatty acids. Together with the block at PDHc and OGDHc, such a mechanism may further restrict substrate flexibility (Fig. 5C).

## Medium-chain triglyceride-enriched diet mitigates metabolic collapse

Maternal milk is rich in medium-chain triglycerides, which enter mitochondria independently of CPT1 (*1*, *56*). We therefore hypothesized that dietary supplementation with MCTs could support β-oxidation and alleviate the emerging energetic crisis. MCT-fed *Slc25a26^KO^* mice displayed improved HW/BW ratio, increased circulating blood triglycerides, reduced cardiomyocyte vacuolization (Fig. 5D and E, fig. S8A to C), and restoration of high-energy metabolites including ATP, UTP, and UDP at P14 (Fig. 5F to H). Citrate levels were increased, likely reflecting higher acetyl-CoA availability in the MCT-fed group, while downstream TCA intermediates beyond αKG remained unchanged (Fig. 5G). Proteomics showed partial normalization with reductions in sarcomeric stress and cardiomyopathy markers and enrichment of cardiac muscle contraction pathways (Fig. 5I and J, fig. S9A to C, data S1).

These improvements suggest that MCT supplementation stabilizes cardiac metabolism by providing an alternative fuel source and, potentially, by enhancing residual lipoylation capacity. Comparing the levels of lipoylated DLAT and DLST between *Slc25a26^KO^* with and without MCT supplementation showed a stable, although mild increase (fig. S9D). Together, our findings demonstrate that dietary MCT can partially restore metabolic homeostasis and cardiac structure in *Slc25a26^KO^*hearts.

## Discussion

The shift toward oxidative metabolism in the early postnatal heart is a tightly coordinated process involving both transcriptional and metabolic reprogramming. Prior studies have implicated SAM levels in regulating chromatin accessibility and epigenetic remodeling during this transition (*29–31*). Our work extends this knowledge by providing the first mechanistically comprehensive characterization of mitoSAM as a critical determinant of cardiac maturation, acting primarily through its role in sustaining protein lipoylation.

MitoSAM contributes to mitochondrial function through two distinct pathways: by supporting mitochondrial methylation reactions or by fueling lipoic acid biosynthesis. Our data show that even a mild reduction in mitoSAM is sufficient to impair lipoylation, selectively blocking oxoacid dehydrogenase activity and creating a metabolic bottleneck at OGDHc that precedes proteome-wide changes or stress marker induction. Importantly, although *Slc25a26* was deleted during late embryogenesis, the decline in mitoSAM levels emerged only after birth, coinciding with the postnatal metabolic shift toward oxidative metabolism. These results highlight lipoic acid biosynthesis as the dominant vulnerability in the postnatally developing heart.

Loss of lipoylation blocks the activity of OGDHc, leading to accumulation of αKG combined with the depletion of aspartate and nucleotide pools. This metabolic imbalance coincides with sustained cardiomyocyte proliferation, suggesting a failure to fully exit the cell cycle and creating a dual bioenergetic and biosynthetic deficit. Interestingly, while previous studies have linked elevated αKG and aspartate availability to enhanced proliferation and cardiac hypertrophy (*16*, *50*, *57*), our findings reveal that αKG accumulation is instead accompanied by reduced aspartate and nucleotide synthesis, consistent with impaired OGDHc activity. Moreover, although αKG has been reported to drive hypomethylation we observe no changes in global 5hmC levels. Together, these results imply that the proliferative phenotype arises from a direct metabolic block at OGDHc rather than from altered methylation.

Notably, the onset of metabolic collapse in *Slc25a26^KO^*hearts coincides with the suckling-to-weaning transition, when cardiac substrate flexibility is critical. This normal developmental shift reduces medium-chain fatty acid availability for both β-oxidation and octanoate-dependent lipoic acid synthesis, thereby compounding the intrinsic lipoylation defect. The rescue of metabolic failure by dietary MCT supplementation highlights the importance of aligning cofactor demand with nutrient availability during this narrow postnatal window.

Our findings thus reveal a previously unrecognized developmental checkpoint in which mitoSAM availability gates the transition from a proliferative to a mature metabolic state. This checkpoint operates through lipoylation-dependent control of TCA cycle flux, linking mitochondrial cofactor supply to nucleotide metabolism and cell cycle exit. This stage-specific vulnerability is supported by a recent study showing that deletion of *Slc25a26* in the adult heart is tolerated unless hearts are challenged with transverse aortic constriction (*58*), emphasizing that the critical requirement for mitoSAM-dependent lipoylation is confined to the period of rapid postnatal growth and metabolic remodeling. Similar developmental windows exist in other organs, such as brain, liver, and kidney (*59–63*) raising the possibility that mitoSAM availability acts as a general regulator of tissue maturation timing in mammalian physiology and disease.

## Acknowledgments

Mass spectrometry analysis was performed by the Clinical Proteomics Mass Spectrometry facility at Karolinska Institutet/Karolinska University Hospital/Science for Life Laboratory. Histological analysis was performed in the Karolinska Institutet Unit for Morphological Phenotype Analysis (FENO). The authors used ChatGPT to improve the readability and language of the manuscript’s first draft. The authors reviewed and edited the suggested changes and take full responsibility for the content of the publication.

## Funding

Swedish Research Council VR2022-01287 (AWr)

Swedish Research Council VR2023-07091 (AWr)

Swedish Research Council VR2023-02388 (AWe)

NovoNordisk Foundation NN0082202 (AWr)

Knut & Alice Wallenberg Foundation KAW2019.0109 (AWr, AWe)

Knut & Alice Wallenberg Foundation KAW2024.0081 (AWr, AWe)

Knut & Alice Wallenberg Foundation KAW2020.0228 (AWe)

Swedish Heart and Lung Foundation 20210498 (AWr, GP, DCA, CF)

Swedish Heart and Lung Foundation 20190310 (LHL)

Swedish Heart and Lung Foundation 20220540 (LHL)

Swedish Heart and Lung Foundation 20230594 (DCA)

Region Stockholm RS2022-0708 (AWr)

Region Stockholm FoUI-955096 (AWe)

Region Stockholm FoUI-987428 (DCA)

Karolinska Institutet consolidator grant 2-190/2022 (AWr)

.Cancerfonden 21 1621 Pj (AWr)

Cancerfonden 25 4595 Pj (AWr)

Promobilia A23059 (DCA)

National Heart Lung and Blood Institute 5F32HL174084-02 (TST)

Cancer Prevention Research Institute of Texas (CPRIT Core Facilities Support) RP240494 (RJD, TPM, RW, and the Children’s Research Institute Metabolomics Facility

The Howard Hughes Medical Institute Investigator Program (RJD)

The National Cancer Institute R35CA220449 (RJD)

## Author contribution

*Conceptualization*: AR, CF, AWr.

*Data curation*: AR, WC, TST, TPM.

*Formal analysis*: AR, WC, AV, LM-W.

*Investigation*: AR, AWi, TST, MM, FAR, DM, AV, YH, DA, RWi, GP.

*Methodology*: AR, TST, MM, AV, YH, DA, RW, TPM, GP.

*Resources*: LHL, DCA, AWe, RJD, CF, AWr.

*Supervision*: RJD, CF, AWr.

*Visualization*: AR, WC, CF.

*Writing original draft*: AR, CF, AWr.

*Reviewing & editing*: all authors.

*Funding*: DCA, LHL, AWe, RJD, AWr.

*Project administration*: AR, CF, AWr.

