## Supplemental information for "MitoSAM-dependent lipoylation controls postnatal heart development via metabolic remodeling"

**The PDF file includes:**

Materials and Methods

Figs. S1 to S9

Tables S1

References 64-83

Materials and Methods

Mouse breeding and tissue collection

*Slc25a26*$^{LoxP/LoxP}$ mice described previously (*27*) were crossed to *Ckmm-Cre* mice (*64*). All mice used in experiments were of C57BL/6N genetic background. Mice were kept in individually ventilated cages at ambient room temperature of 22 – 24°C, had ad libitum access to food and water, and were maintained with 12 h:12 h light:dark cycle. Groups included male and female animals. Medium chain triglycerides diet (sniff Spezialdiäten GmbH) consisting of 56 kJ% fat (containing 30% hydrogenated coconut oil, C8 content 1.77%), 27 kJ% carbohydrates 17 kJ% was provided in a Petri dish, placed on a bottom of the cage, when pups were 11 days old. It was replaced every 2 days. All the time, the breeding pair had ad libitum access to normal chow placed on a wire rack above the cage bedding. Genotyping primers for *Slc25a26*$^{LoxP}$ alleles:

*Slc25a26*$^{LoxP}$ primers: Fw 5′-ACA GCT GAG TGC AGT AGA GAC C-3′; Rv 5′-CCA TAA GAG ATA AGG CAC AGA GG-3′. *Cre* primers: Fw 5′-CAC GAC CAA GTG ACA GCA AT-3′; Rv 5′-AGA GAC GGA AAT CCA TCG CT-3′.

Blood triglyceride levels were determined with Accutrend Plus device (Roche Diagnostics) using Accutrend triglycerides test strips (Roche Diagnostics). Following euthanasia by cervical dislocation or decapitation for animals younger than P10, dissected organs were snap-frozen in liquid nitrogen and used further for proteome, metabolome analyses, gDNA, mRNA, and protein extraction. Crude mitochondria isolation for respiratory chain enzyme activities assay and *ex vivo* ATP production rates were done from freshly dissected organs. Tissues for histological analyses were fixed in 10% (v/v) neutral buffered formalin overnight at 4°C; the next day, tissues were transferred to 70% (v/v) ethanol and stored at 4°C until further use. All animal experiments were approved by the local animal welfare committee. All procedures were conducted in accordance with institutional, national, and European recommendations and followed the guidelines of the Federation of European Laboratory Animals Associations (FELASA).

Transthoracic echocardiography

Cardiac function was non-invasively monitored by transthoracic echocardiography using a high-resolution imaging system (Philips HDI 5000) with a CL-15 7 scanning head, as described previously (*65*, *66*). Briefly, 12-day-old mice were anesthetized with isoflurane (2%) and laid on a heating pad (37°C). Chest hair was removed using depilatory cream. After echocardiography image acquisition, the mice were maintained on a heating pad (37°C) without anesthesia until their recovery. M-mode imaging of the left ventricle short axis was analyzed, and cardiac contractility was expressed as % of left ventricular fractional shortening using the following formula: LVd-LVs/LVd *100 where LVd and LVs stand for left ventricle diastole and systole respectively. Heart rate (HR), posterior wall (PW), and interventricular septum (IVS) thickness were also measured.

EdU administration

Click-iT EdU Cell Proliferation Kit (Thermo Fisher Scientific) was used according to the manufacturer's instructions to detect proliferating cells in vivo. Briefly, mice were injected intraperitoneally with EdU dissolved in DMSO (dose 20 mg per kg body weight) and kept alive for 4 h for in vivo labeling. Afterwards, mice were euthanized as usual; heart and duodenum (positive control of EdU incorporation) were dissected and fixed overnight at 4°C in 10% normal buffered formalin. Paraffin embedding and sectioning (10 µm) were done according to standard procedures. Each slide included heart and duodenum sections. Sections were deparaffinized in xylene and rehydrated in gradual alcohol concentrations and water. Click-iT chemistry reaction was performed according to the manufacturer's instructions. After click-iT chemistry reaction sections were processed further for immunofluorescent staining.

Immunofluorescent staining of heart sections

Formalin-fixed hearts were paraffin-embedded and sectioned (10 µm). Deparaffinization was done in xylene followed by the rehydration in gradual alcohol concentrations and water. Following EdU click-iT chemistry reaction, sections were blocked in 10% normal goat serum in PBS for 30 min and stained with cardiac troponin T antibody (Invitrogen #MA5-12960) overnight at 4°C. The next day sections were washed and stained with anti-mouse IgG secondary antibody (Invitrogen #A11017) for 2 h at room temperature, followed by washes and DAPI-staining of nuclei for 5 min at room temperature. Wheat-germ-agglutin conjugated to fluorophore (Invitrogen) was applied for 2 h at room temperature.

Hematoxylin and eosin staining of heart sections

Formalin-fixed hearts were paraffin-embedded, sectioned (10 µm), deparaffinized and rehydrated according to standard procedures. Sections were stained with Mayer's hematoxylin and eosin. Embedding, sectioning, staining, and image acquisition were done by the Karolinska Institutet Unit for Morphological Phenotype Analysis (FENO).

Crude mitochondria isolation

Dissected hearts were washed in ice-cold 0.01 M PBS and cut into small pieces. The pieces were transferred to an isolation buffer (100 mM sucrose, 50 mM KCl, 1 mM EDTA, 20 mM TES 0.2% (w/v) BSA, pH 7.2, supplemented with subtilisin A from *Bacillus licheniformis*) and homogenized with 20 strokes at 1000 rpm (Schuett homgen$^{plus}$). Homogenates were differentially centrifuged at 800 *rcf* and 8500 *rcf*. Mitochondrial fractions from hearts were resuspended in the isolation buffer without BSA. Protein concentration was determined with Qubit™ Protein Assay (Thermo Fisher Scientific).

Blue Native Electrophoresis and in-gel activity assay

Ten micrograms of mitochondria were lysed with 4% digitonin to resolve supercomplexes. BN-PAGE was performed in NativePAGE Novex Bis-Tris Gel System (Life Technologies) according to the manufacturer's instructions. In-gel activity assays for complex I and complex IV were performed as previously described (*67*).

Isolated respiratory chain enzyme activities

Electron transport chain enzyme complex activities and citrate synthase activity from isolated mouse heart mitochondria were measured as described previously (*68*). All ETC activities were normalized to the citrate synthase activity in the mitochondrial suspension.

Ex vivo ATP production rates

Ex vivo ATP production rates were measured as described previously (*68*). Freshly isolated mitochondria from mouse heart were immediately resuspended in a buffer (250 mM sucrose, 15 mM $KH_2PO_4$, 2 mM magnesium acetate, 0.5 mM EDTA and 0.5 g/L BSA pH 7.2) 10 µL of resuspended mitochondria were thereafter added to p96 wells each containing a luciferase/luciferin-based ATP monitoring reagent (BioThema SL) and a different combination of substrates: 15 mM glutamate + 15 mM succinate; 30 mM glutamate + 20 mM malate; 30 mM pyruvate + 10 mM malate; 20 mM α-ketoglutarate; 20 mM succinate + 1 mg/L rotenone; 20 µM decanoyl-L- carnitine + 1 mM malate; 10 µM lauryl-L- carnitine +1 mM malate; 5 µM myristoyl-L-carnitine + 1 mM malate; 5 µM palmitoyl-L-carnitine +1 mM malate. Reaction was initiated by injecting 15 µL of ADP at a final concentration of 0.6 mM. Luminescence increase was followed during 7 min in a Tecan SPARK microplate reader at a constant temperature of 28 °C. After, an ATP internal standard (final concentration 0.5 µM) was injected in each well and luminescence increase was used to convert counts to ATP μmols. Each substrate was measured in technical triplicates. Obtained results were expressed as Units (μmol ATP/min) normalized by units of citrate synthase. Content in microplate wells at a final volume of 250 μL: 150 mM sucrose, 15 mM $K_2HPO_4$, 2 mM $MgAc_2$, 0.5 mM EDTA, firefly luciferase, 0.1 g/L D-luciferin, 0.004 g/L L-luciferin, 1 g/L BSA, 1 μM $Na_2P_2O_7$, 0.6 mM ADP, 1 mM AMP, 0.2 μM DAPP and 0.5 μM ATP (added as internal standard at the end of the measurements).

$CoQ_9$ and $CoQ_{10}$ levels measurement

$CoQ_9$ and $CoQ_{10}$ levels were measured as described previously (*69*). Briefly, isolated mitochondria were resuspended in 200 μL of 1 mM $CuSO_4$, followed by the addition of 200 μL ethanol. Samples were ultrasonicated at high power for 5 min in an ice-cold water bath (Bioruptor®, Diagenode). After adding hexane (400 μL), the samples were vortexed and centrifuged at 2500 *rcf* for 5 min at 4°C. The hexane-soluble upper fraction was collected and dried using a vacuum concentrator (Eppendorf). $CoQ_9$ and $CoQ_{10}$ levels were normalized to protein content.

SAM and SAH levels

SAM and SAH levels were measured as previously described (*27*). In brief, metabolites were extracted from mitochondrial pellets or total homogenates with ice-cold 80% (v/v) methanol spiked with labeled internal standards SAM[$^2H_3$] and SAH[$^2H_4$], followed by incubation on ice. The extracts were analyzed using a Waters (XEVO TQ-XS, Milford, US) instrument. Data acquisition and peak integration were performed with MassLynx 4.1 software (Waters). SAM and SAH levels were normalized to protein content.

Untargeted metabolomics of mouse hearts

Samples were stored at −80°C until processing. Metabolites were homogenized in pre-cooled (4°C) 80% acetonitrile (ACN) using a Qiagen TissueLyser III system. Homogenates were centrifuged at 18,000 *rcf* for 15 min at 4°C, and the supernatant was transferred to a new tube. Residual protein content was quantified using the Pierce™ BCA Protein Assay (Thermo Fisher Scientific), and all samples were normalized to 0.1 μg protein/μL in 80% ACN.

Metabolites were analyzed by liquid chromatography-high resolution tandem mass spectrometry as reported previously (*70*, *71*). Briefly, extracts were loaded onto a QExactive HF-X mass spectrometer coupled to a Vanquish Flex liquid chromatography system (Thermo

Scientific, Bremen, Germany). Metabolites were chromatographically resolved with a Millipore SeQuant ZIC-pHILIC column and a binary gradient of 10 mM ammonium acetate in water, pH 9.8 and ACN. Precursor and product ion spectra of metabolites were acquired in both positive and negative polarities. Metabolites were quantified with Compound Discoverer 3.3 searching against an in-house spectral library of previously validated metabolites. Principal component analyses, pathway enrichment and statistical analyses were done using MetaboAnalyst 6.0 (*72*).

RNA isolation and quantitative PCR with reverse transcription

Hearts were stored at −80°C until processing. Tissue was homogenized with Fastprep-24 tissue homogenizer (MP Biomedicals) with 2 x 30 s at 6000 rpm in TRIzol™ (Thermo Fisher Scientific). mRNA was extracted using a standard TRIzol™/chloroform extraction method. RNA concentrations were measured with Nanodrop ND-1000 (Thermo Fisher Scientific). Isolated RNA was treated with DNase (DNA-free Kit, Ambion) and subsequently reverse-transcribed with the High-Capacity cDNA Reverse Transcription Kit (Applied Biosystems). Quantitative reverse transcription PCR was performed in a QuantStudio 6 system using TaqMan probes and TaqMan Universal Master Mix II, with uracil-N-glycosylase (UNG) (Thermo Fisher Scientific). Normalization was done to *Hprt*, *Actb*, or *18S*. The relative expression of mRNAs was determined with a comparative method ($2^{-\Delta\Delta Ct}$). The list of TaqMan probes can be found in the table S1.

SDS-PAGE and immunoblot analysis

Samples were stored at −80°C until processing. Heart and quadriceps tissue were homogenized with Qiagen TissueLyser III system in T-PER™ Tissue Protein Extraction Reagent (Thermo Scientific), containing protease inhibitor cocktail (Roche) and phosphatase inhibitor cocktail (PhosSTOP, Roche). Protein concentration was determined with Pierce™ BCA Protein Assay (Thermo Fisher Scientific). Protein lysates (20-30 µg) were resuspended in the NuPAGE™ LDS Sample Buffer (Thermo Fisher Scientific), incubated at 70°C for 10 min and loaded onto precast Bolt™ 4% – 12% or 12% Bolt™ Bis-Tris gels (Thermo Fisher Scientific) in an XCell SureLock electrophoresis system (Thermo Fisher Scientific). Following electrophoresis, proteins were transferred to nitrocellulose or PVDF membranes, blocked in EveryBlot Blocking Buffer (Bio-Rad) for 5 min at room temperature or 5% fat-free milk in TBST for 1 h and immunoblotted with primary antibodies. Incubation with primary antibodies was done overnight, at 4°C, and secondary antibodies anti-mouse IgG (Cytiva, #NA9310) or anti-rabbit IgG (Cytiva, #NA9340) conjugated to HPR were applied for 1 h at room temperature. Clarity Western ECL Substrate (Bio-

Rad) was applied. Immunoblot images were acquired with ChemiDoc™ XRS+ (Bio-Rad). Stripping of nitrocellulose membranes, if needed, was done with Restore™ Western Blot Stripping Buffer (Thermo Fisher Scientific). The following primary antibodies were used: rodent OXPHOS cocktail (Abcam #ab110413), Vinculin (CST #4650), HSC70 (Santa Cruz #sc-729), α-Tubulin (Millipore #CP06), GAPDH (Abcam #ab8245), TOMM20 (Sigma-Aldrich #HPA011562), DLAT (Proteintech #13426-1-AP), DLST (Sigma-Aldrich #HPA003010), OGDHE1 (Sigma-Aldrich #HPA020347), Citrate synthase (Santa Cruz #sc-390693), Aconitase-2 (Abcam #ab110321), lipoic acid (Millipore #437695), ACC (CST #3662), phospho-ACC (Ser79) (CST #3661).

Dot blot

Hearts were stored at −80°C until processing. Genomic DNA was isolated with the high-salt chloroform method (*73*). The concentration was determined with Nanodrop ND-1000 (Thermo Fisher Scientific), and all samples were diluted to 50 ng/mL. 100 mL of gDNA was mixed with an equal volume of DNA denaturing buffer (200 mM NaOH, 20 mM EDTA) and denatured at 95°C for 10 min followed by immediate cooling on ice for 5 min. Samples were mixed with 200 mL 20x saline-sodium buffer (3 M NaCl, 0.3 M sodium citrate pH 7.4) and brought up to 500 mL with nuclease-free water. A 2-fold serial dilution was prepared. A positively charge Hybond™-N+ nylon membrane (Cytiva) was prewetted in 2x saline-sodium buffer for 10 min and assembled in a dot blot apparatus (Carl Roth). 100 mL of each dilution was loaded into individual wells and allowed to filter through the membrane under a gentle vacuum. The dot blot apparatus was disassembled; the membrane was air-dried for 5 min and UV cross-linked for 30 s with 120,000 mJ/cm2. The membrane was blocked in EveryBlot Blocking Buffer (Bio-Rad) for 5 min at room temperature and incubated with anti-5hmC antibody (Active Motif #39769-AF) for 1 h at room temperature. Afterwards, the membrane was washed 3 times TBST buffer, followed by the incubation with secondary anti-rabbit antibody Cytiva, #NA9340) conjugated to HPR for 1 h at room temperature. Clarity Western ECL Substrate (Bio-Rad) was applied. Immunoblot images were acquired with ChemiDoc™ XRS+ (Bio-Rad).

Sample preparation for proteomics

Hearts were stored at −80°C until processing. Samples were cryogenically pulverized using the Covaris CryoPREP system (Covaris, Woburn, MA, USA). 200 mL of lysis buffer (4% SDS, 50 mM HEPES pH=7.6, 1 mM DTT) was added, samples were heated to 95°C, and sonicated with Ultrasonic Processor UP200St (Hielscher Ultrasonics, Teltow, Germany). Lysates were centrifuged, and supernatants collected for digestion. For each sample, 200 mg of protein was reduced for 30 min in lysis buffer at 37°C and subsequently alkylated with 10 mM chloroacetamide (CAA) for 10 min at room temperature in the dark. Protein cleanup and digestion were performed using the SP3 method (*74*, *75*). Briefly, proteins were bound to SeraMag SP3 bead mix (40 µL) in 70% ACN, washed twice with 70% ethanol and once with 100% ACN. The beads-protein mixture was reconstituted in 100 µL digestion buffer (50 mM HEPES pH 7.6, 10% ACN, 10 mM $CaCl_2$ and 0.4 µg Rapizyme) and incubated overnight at 37°C. The following day, the peptides were cleaned by adding 100% ACN and digested overnight with Rapizyme Trypsin (MS grade, Waters) in the digestion buffer (50 mM HEPES buffer pH 7.6, 10% ACN and 10 mM $CaCl_2$). Peptides were recovered by >95% ACN precipitation, washed, and resuspended in 0.1% formic acid for LC–MS injection.

LC-ESI-MS/MS (DIA, timsTOF HT)

500 mg of peptides was loaded onto Evotip PURE (EvoSep). Separation was performed using an Evosep One system with a 30 samples/day gradient on a PepSep FIFTEEN column (15 cm × 150 µm, 1.5 µm; Bruker) at 40°C. Mobile phases were: A = 0.1% FA in water, B = 0.1% FA in acetonitrile. Eluting peptides were analyzed on a timsTOF HT mass spectrometer (Bruker) equipped with a CaptiveSpray source. Data were acquired in dia-PASEF mode with an m/z range of 300–1300 and ion mobility 0.6–1.6 $1/k_0$. Isolation windows were optimized with Py-DiaID (*76*). Collision energies were ramped from 15–70 eV, with a total cycle time of 1.8 s.

The separation column was a PepSep FIFTEEN 15 cm x 150 µm x 1.5 µm (Bruker 1893474) connected to a 20 µM ZDV Sprayer (Bruker 1865710) at 40°C. Mobile phase A was 0.1% FA in MQ and B 0.1% FA in ACN. Online LC-MS was performed using a timsTOF HT mass spectrometer (Bruker) using the CaptiveSpray source, capillary voltage 1500 V, dry gas flow of 3 L/min, dry gas temperature at 180°C. Collision energy was set as 20 eV for 1/k0 0.60 Vs/cm and 59 eV for 1/k0 1.60 Vs/$cm^2$. Data was acquired using Timscontrol v 6.0.6 and Compass HyStar 6.3.1.8. The above conditions were used for data independent acquisition (DIA) diaPASEF (parallel accumulation-serial fragmentation). For dia-PASEF analysis; precursor ions were selected in the range of 300-1300 m/z and 0.6-1.6 1/k0 IM. Optimized isolation windows based on the Bruker human library with the assistance of the Py-DiaId Python package (*76*) were implemented. Collision energies for fragmentation were ramped from 15 to 70 eV. Parallel accumulation and serial fragmentation were performed with a cycle time of 1.8 s.

Proteomic data analysis (DIA)

Raw DIA files were processed in Spectronaut (Biognosys) using the DirectDIA workflow for label-free quantification. A project-specific spectral library was generated directly from the DIA data. Search parameters: 1% false discovery rate (FDR) at peptide and protein level, up to 2 missed cleavages, enzyme specificity = trypsin (*77*). Searches were performed against the UniProt mouse proteome (ID: 10088). Protein quantification was based on the top 3 peptides per major and top 6 per minor group, using only unique peptides mapped to a single gene ID.

Downstream statistical analysis of proteomics data

MS2-based protein intensity values exported from Spectronaut (DIA datasets), Proteome Discoverer (TMT datasets), or MaxQuant (LFQ datasets) were processed in R (v4.5.1). Protein-level expression analysis followed the standard DEP2 workflow (*78*). Briefly, data were normalized using normalize_vsn() (variance-stabilizing transformation), and differential abundance was tested using test_diff(). Proteins were considered significantly altered at adjusted $p < 0.05$ and $|\log_2 FC| > 0.58$, unless otherwise indicated. Functional enrichment analysis was performed with clusterProfiler (*79*), applying Gene Ontology (GO), KEGG pathway and Reactome terms. Enrichment results were visualized using clusterProfiler plotting functions and ggplot2.

Sample preparation for redox proteomics

Hearts were stored at −80°C until processing.  Samples were thawed on ice, minced, and transferred to tubes containing 400 µm LoBind silica beads with 100 µL of lysis buffer (4 M urea, 0.1% ProteaseMAX (Promega) and 50 mM NaCl in 100 mM Tris-HCl, pH 8.5). Samples were homogenized with a Disruptor Genie (2800 rpm, 2 min cycles, repeated ×5 with cooling) and centrifuged (14,000 *rcf*, 10 min, 4°C). Supernatants were collected, sonicated (40 s, 20% amplitude, 2 s on/2 s off), and protein concentrations were measured with Pierce™ BCA Protein Assay (Thermo Fisher).

For redox labeling, 50 µg aliquots were adjusted to 20 mM EPPS buffer pH 8.5 and alkylated with 50 mM iodoacetamide (IAA) for 1 h at 25°C in the dark. Proteins were precipitated (chloroform/methanol), resuspended in 8 M urea, sonicated, and diluted in EPPS buffer pH 7.2. Oxidized cysteines were reduced with 0.5 mM DTT (45 min, 25°C) and re-alkylated with 100 mM N-ethylmaleimide (2 h, 25°C, dark). Proteins were reprecipitated, resuspended, and digested with trypsin (1 μg, 6 h, 25°C). Peptides were cleaned on a HyperSep C18 plate (Thermo Fisher) and dried by vacuum centrifugation.

LC–MS/MS (DDA, Orbitrap Q Exactive HF)

Peptides (~2 μg) were injected onto an EASY-Spray C18 column (50 cm, Thermo Fisher) coupled to an Ultimate 3000 nanoUPLC (Thermo Fisher). Separation was achieved with a 90 min gradient (4–26% B in 90 min, 26–95% B in 5 min; flow 300 nL/min). Mobile phase A: 0.1% FA in water; B: 0.1% FA in 98% acetonitrile ACN.

MS acquisition was performed on a Q Exactive HF (Thermo Fisher) with full MS scans from m/z 375–1500 at R=120,000, AGC target $5\times10^6$, max injection time 80 ms. Data-dependent MS/MS of the top 18 precursors (charge 2+–7+) was performed with HCD (NCE 33%), R=60,000, AGC target $2\times10^5$, max IT 54 ms, isolation width 1.4 Th, and 45 s dynamic exclusion.

Redox proteomics downstream data analysis

MT datasets: Processed in Proteome Discoverer v3.0 (Thermo Fisher) with Mascot v2.5.1 search engine. Database: SwissProt mouse proteome. Parameters: 2 missed cleavages, precursor mass tolerance 10 ppm, fragment 0.02 Da. Fixed modifications: carbamidomethylation and N-ethylmaleimide (Cys), TMTpro (Lys, N-terminus). Variable: Met oxidation, Asn/Gln deamidation. FDR ≤5% at peptide level using Percolator. Downstream statistical analyses were performed in R using limma (*80*).

Single-cell RNA-seq data analysis

Single-cell RNA-seq data from murine heart were obtained from the Tabula Muris dataset (*28*) in Seurat object format. Data processing and visualization were carried out in R (version 4.5.1) using the Seuratpackage (*81*). Standard preprocessing steps included normalization with SCTransform(), dimensionality reduction with RunPCA(), RunUMAP(), and RunTSNE(), followed by neighborhood graph construction with FindNeighbours() and clustering using FindClusters(). Cell-type annotation of resulting clusters was performed with the clustermole package https://igordot.github.io/clustermole/. Visualization and plotting were carried out with Seurat functions and ggplot2.

Immunofluorescent image capture and quantification

Immunofluorescent-stained sections were imaged with a confocal laser-scanning microscope (Leica Stellaris 5 X, Leica Microsystems). *Z*-stack images were acquired following the Nyquist sampling in sequential mode. Leica Application Suite X software (Leica Microsystems) and Fiji (ImageJ) were used further to set up uniform contrast and brightness. For cardiomyocyte cross-sectional area 900 cardiomyocytes per heart section were manually traced and their area was measured in Fiji. The number of $EdU^+$ cardiomyocytes per square millimeter of heart was assessed by counting the number of $EdU^+$ cardiomyocytes in a region of interest, defined in Fiji (ImageJ) and dividing it by the area of region of interest in millimeters squared.

Quantification, statistical analysis and visualization

The number of mice analyzed per experiment is indicated in corresponding figure legends. Statistical significance between two groups was assessed using Student's *t*-test, one-way and two-way ANOVA were applied for multiple comparisons. Analyses were carried out in Prism 10 (GraphPad Software). The number of replicates and specific statistical test are detailed in each figure legend and methods. Unless stated otherwise, values represent mean ± SD. No statistical method was used to predetermine sample size. Figure preparation was done using Prism 10 (GraphPad Software) and Adobe Illustrator 2024 (Adobe).

Data availability

The proteomics data generated in this study have been deposited in the ProteomeXchange via the jPOST repository (*82*).

Raw redox proteomics data are deposited at ProteomeXchange via PRIDE (*83*)

Datasets used in this study are MitoXplorer v2, FASTA files from UniProt (*M. musculus* in September 2018). Source data are provided with this paper.

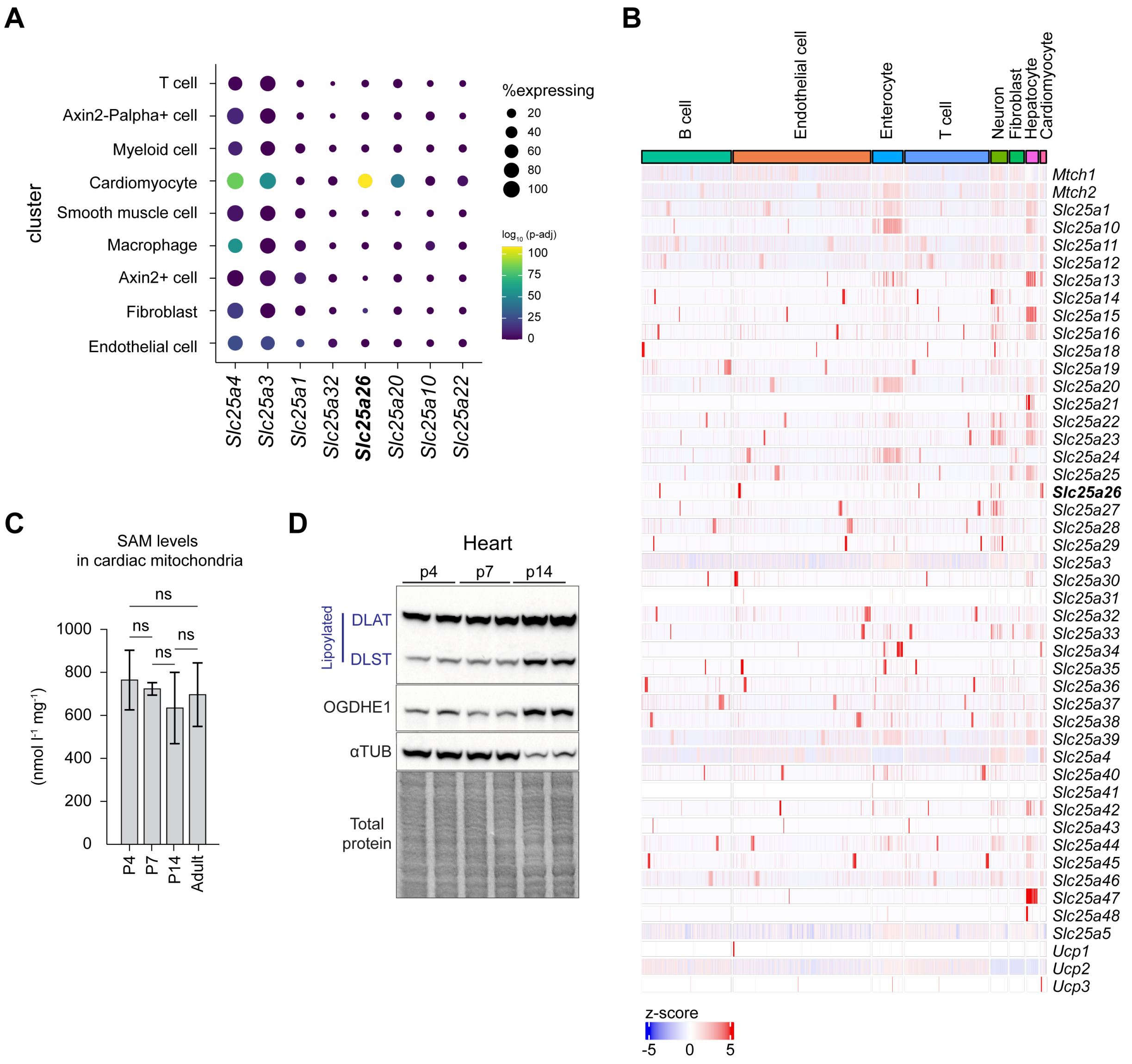


**Fig. S1.** ***Slc25a26*** **expression and lipoylation are maintained during postnatal cardiomyocyte maturation.**

(**A**) Dot plot of selected SLC25 family members in indicated clusters. Data were obtained from a publicly available single-cell dataset (28). The size of the dots indicates the proportion of cells expressing the indicated gene relative to the cluster. Color indicates the significance of the differential expression of the indicated gene. (**B**) Heat map showing all SLC25A family members' expression across the indicated lineages. (**C**) Targeted metabolic analysis for mitoSAM levels at indicated time points in Ctrl cardiac mitochondria (n=2 – 8 per time point). (**D**) Immunoblot of

Ctrl heart lysates at indicated time points, with indicated antibodies; α-tubulin (αTUB) was used as the loading control. Data in (C) presented as mean ± SD, one-way ANOVA, with Tukey's multiple comparisons test; ns = not significant.

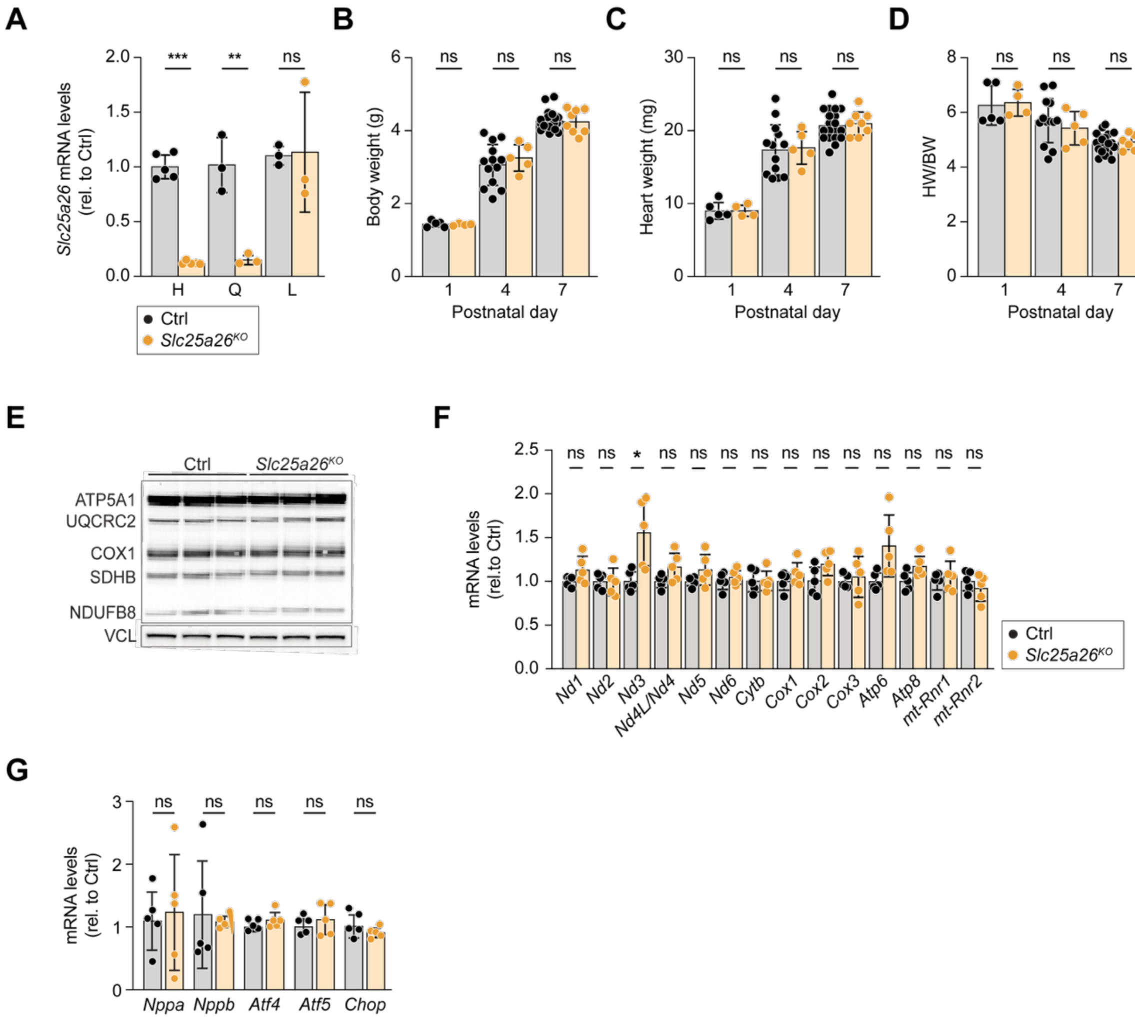


**Fig. S2. *Slc25a26$^{KO}$* mice do not show mitochondrial gene expression defect at P7.**

(**A**) Transcript levels of *Slc25a26* in heart (H), quadriceps (Q) and liver (L) in Ctrl and *Slc25a26$^{KO}$* mice, qRT-PCR (n=5 for heart, n=3 for quadriceps and liver per genotype). (**B**) Body weight (BW) of Ctrl and *Slc25a26$^{KO}$* mice at indicated time points (n=4 – 19 per genotype and time point). (**C**) Heart weight (HW) of Ctrl and *Slc25a26$^{KO}$* mice at the indicated time points (n=4 – 19 per genotype and time point). (**D**) HW normalized to BW of Ctrl and *Slc25a26$^{KO}$* mice at the indicated time points (n=4 – 19 per genotype and time point). (**E**) Immunoblot of Ctrl and *Slc25a26$^{KO}$* heart lysates for indicated ETC complexes subunits at P7. Vinculin (VCL) was used as a loading control (n=3 per genotype). (**F**) Transcript levels of indicated mtDNA-encoded genes in Ctrl and *Slc25a26$^{KO}$* hearts at P7, qRT-PCR (n=5 per genotype). (**G**) Transcript levels of indicated cardiomyopathy and integrated stress-response markers in Ctrl and *Slc25a26*$^{KO}$ hearts at P7, qRT-PCR (n=5 per genotype). Data in (A – D, F and G) presented as mean ± SD, each dot represents an individual mouse, two-tailed unpaired Student's *t*-test, ns = not significant, *$p<0.05$; **$p<0.01$; ***$p<0.001$.

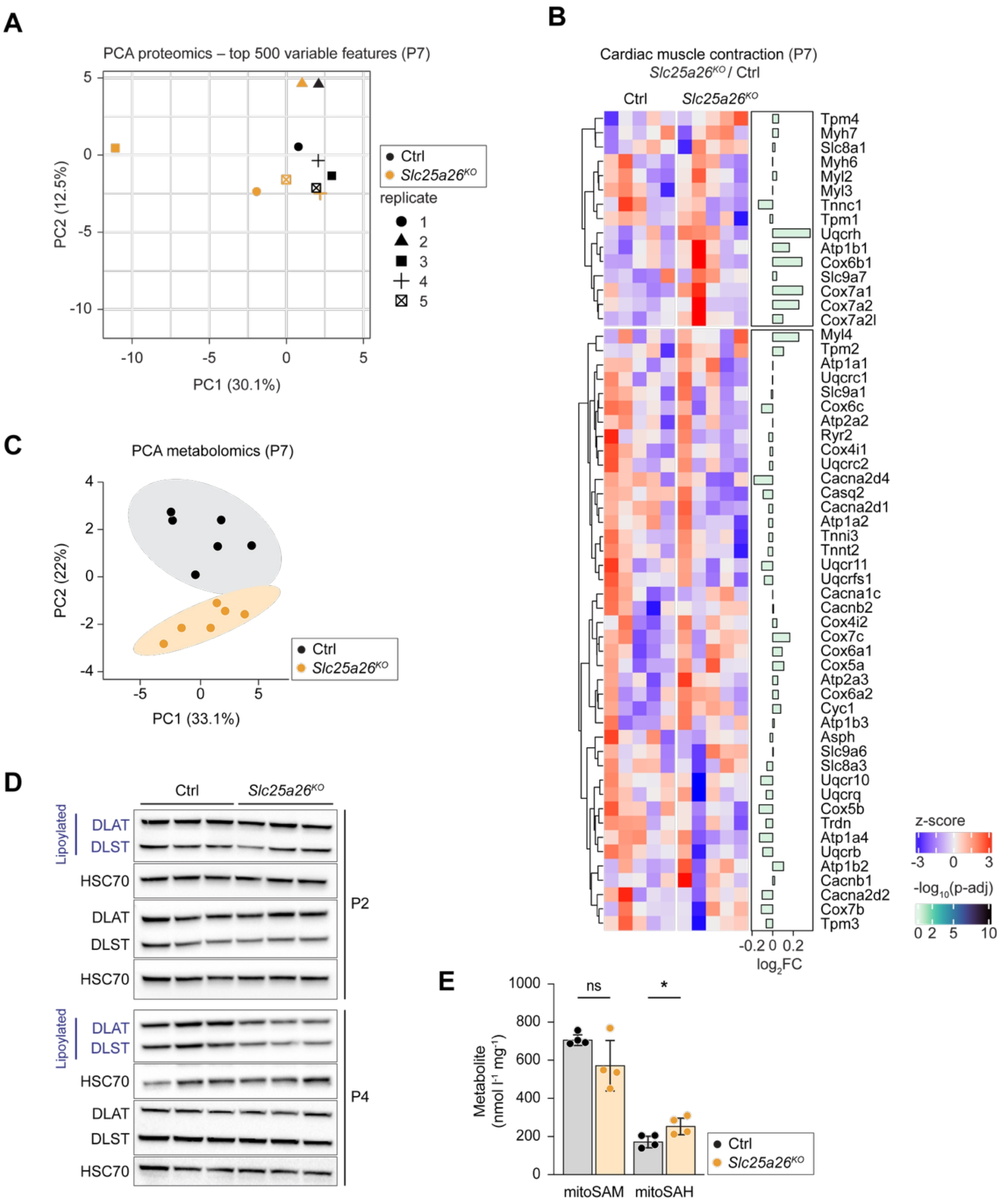


**Fig. S3. *Slc25a26$^{KO}$* mice show progressive lipoylation loss from P4.**

(**A**) Principal component analysis (PCA) of LC-MS/MS label-free whole heart proteomic analysis from Ctrl and *Slc25a26$^{KO}$* mice at P7 (n=5 per genotype). (**B**) Heat map with relative z-score of heart proteomics data, showing proteins in cardiac muscle contraction KEGG pathway,

*Slc25a26*$^{KO}$ hearts compared with Ctrl at P7. (**C**) PCA plot of untargeted metabolomic data in Ctrl and *Slc25a26*$^{KO}$ hearts at P7 (n=6 per genotype). (**D**) Immunoblot of Ctrl and *Slc25a26*$^{KO}$ heart lysates, depicting lipoylated and apo-forms of DLAT and DLST at P2 and P4; HSC70 was used as the loading control (n=3 per genotype). (**E**) Targeted metabolic analysis for mitoSAM and mitoSAH levels in Ctrl and *Slc25a26*$^{KO}$ cardiac mitochondria (n=4 per genotype). Data in (E) presented as mean ± SD, each dot represents an individual mouse, two-tailed unpaired Student's *t*-test, ns = not significant, *$p<0.05$. Statistical analysis for (B) was performed using a linear model and moderated *t*-statistics. *P*-values were adjusted for multiple testing (Benjamini-Hochberg).

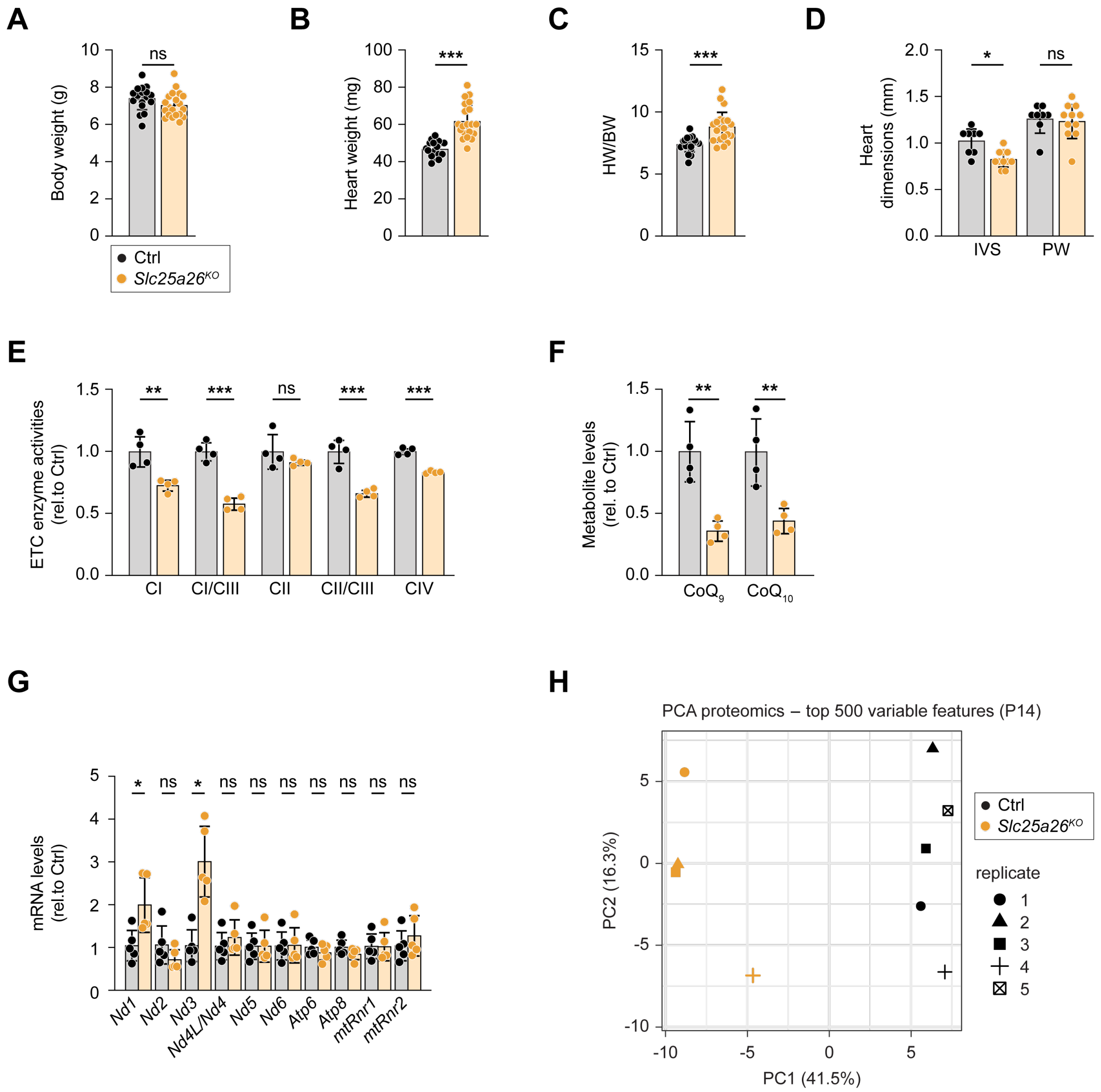


**Fig. S4. *Slc25a26$^{KO}$* mice present with cardiomyopathy at P14.**

(**A**) Body weight (BW) of Ctrl and *Slc25a26$^{KO}$* mice at P14 (n=18 – 20 per genotype). (**B**) Heart weight (HW) of Ctrl and *Slc25a26$^{KO}$* mice at P14 (n=18 – 20 per genotype). (**C**) HW normalized to BW of Ctrl and *Slc25a26$^{KO}$* mice at P14 (n=18 – 20 per genotype). (**D**) Echocardiographic analysis of Ctrl (n=8) and *Slc25a26$^{KO}$* (n=11) mice at P12, dimensions of the intraventricular septum (IVS) and posterior wall (PW). (**E**) Relative isolated electron transport chain (ETC) enzyme activities in Ctrl and *Slc25a26$^{KO}$* cardiac mitochondria (n=4 per genotype) at P14. (**F**) Targeted metabolic analysis for $CoQ_9$ and $CoQ_{10}$ levels in Ctrl and *Slc25a26$^{KO}$* cardiac mitochondria at P14 (n=4 per genotype). (**G**) Transcript levels of indicated mtDNA-encoded genes in Ctrl and *Slc25a26*$^{KO}$ hearts at P14, qRT-PCR (n=5 per genotype). (**H**) PCA plot of LC-MS/MS label-free whole heart proteomics, Ctrl and *Slc25a26*$^{KO}$ at P14 (n=5 for Ctrl, n=4 for *Slc25a26*$^{KO}$). Data in (A – G) presented as mean ± SD, each dot represents an individual mouse, two-tailed unpaired Student's *t*-test, ns = not significant, *$p<0.05$; **$p<0.01$; ***$p<0.001$.

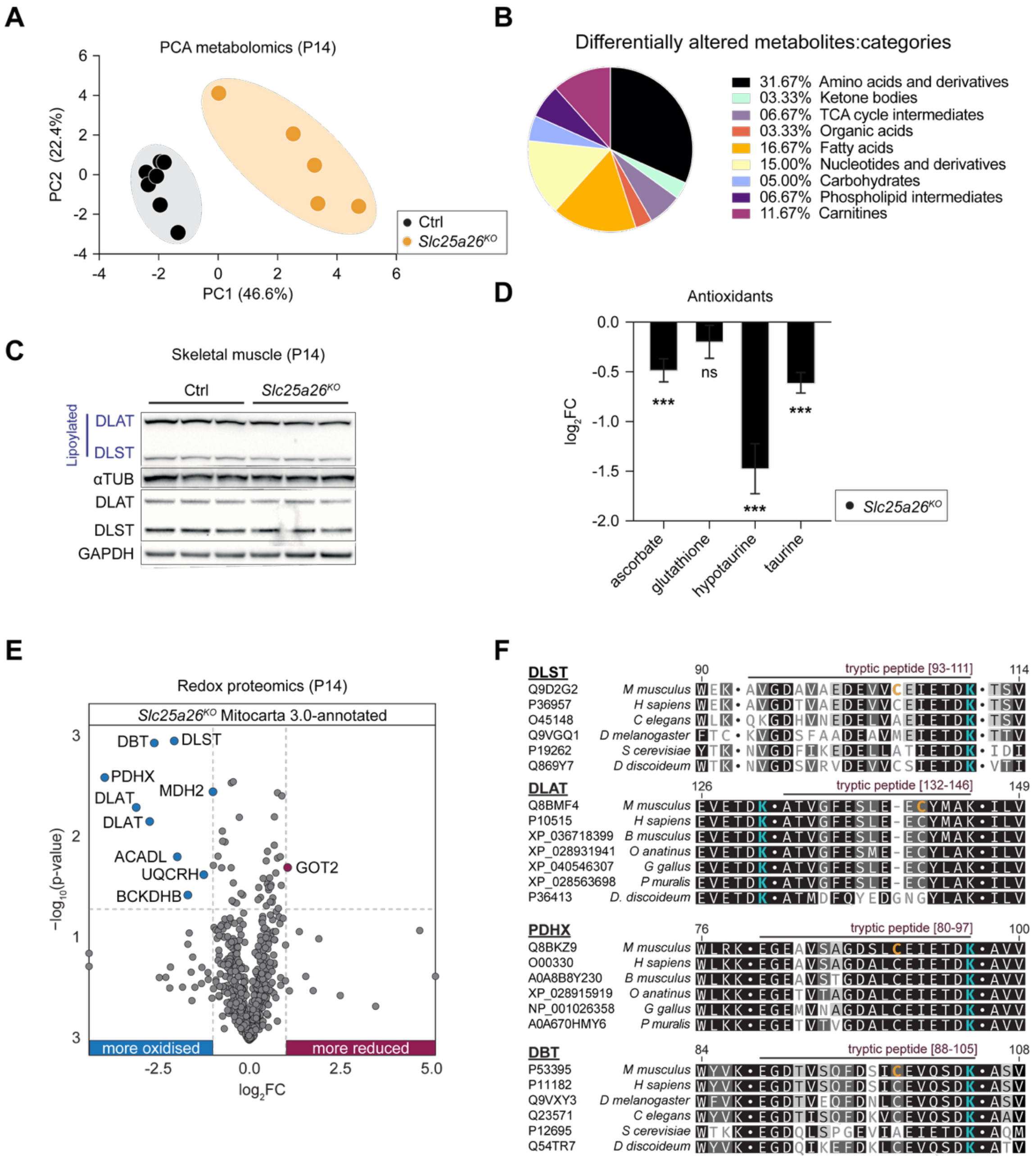


**Fig. S5. *Slc25a26$^{KO}$* deficiency leads to lipoylation loss, altering cysteine oxidation of 2-oxoacid dehydrogenases at P14.**

(**A**) PCA plot of untargeted metabolomic data in Ctrl and *Slc25a26$^{KO}$* hearts at P14 (n=7 for Ctrl, n=5 for *Slc25a26$^{KO}$*). (**B**) Categories of significantly altered metabolites in *Slc25a26$^{KO}$* hearts at P14. (**C**) Immunoblot of Ctrl and *Slc25a26$^{KO}$* quadriceps lysates, depicting lipoylated and apo-forms of DLAT and DLST at P14, α-tubulin (αTUB) and GAPDH were used as the loading controls (n=3 per genotype). (**D**) $Log_2FC$ of antioxidants detected in the untargeted metabolomie of *Slc25a26*$^{KO}$ hearts compared with Ctrl at P14. (**E**) Volcano plot of the mitochondrial redox proteome from *Slc25a26*$^{KO}$ (n=4) vs Ctrl (n=4) hearts at P14. Peptides containing differentially changed cysteines (FC, adjusted $p$-value <0.05) are shown in purple (oxidized) and blue (reduced). (**F**) Protein sequence alignments of the 2-oxoacid dehydrogenase E2 subunits DLST, DLAT, pyruvate dehydrogenase protein X (PDHX), and dihydrolipoamide branched-chain transacylase (DBT). Mouse (*Mus musculus*) was used to align orthologs from humans (*Homo sapiens*), roundworm (*Caenorhabditis elegans*), fruit fly (*Drosophila melanogaster*), brewer's yeast (*Saccharomyces cerevisiae*), amoebae (*Dictyostelium discoideum*), blue whale (*Balaenoptera musculus*), platypus (*Ornithorhynchus anatinus*), chicken, (*Gallus gallus*), and the common wall lizard (*Podarcis muralis*). Differentially oxidized cysteines in the mouse sequences are indicated in orange. Lipoylated lysines are shown in blue. Data in (D) presented as $log_2FC \pm$ s.e.m, two-tailed unpaired Student's $t$-test; ns not significant, ***$p$<0.001. Significance analysis for (E) was performed using a linear model, with moderated $t$-statistics and multiple-testing correction (Benjamini-Hochberg).

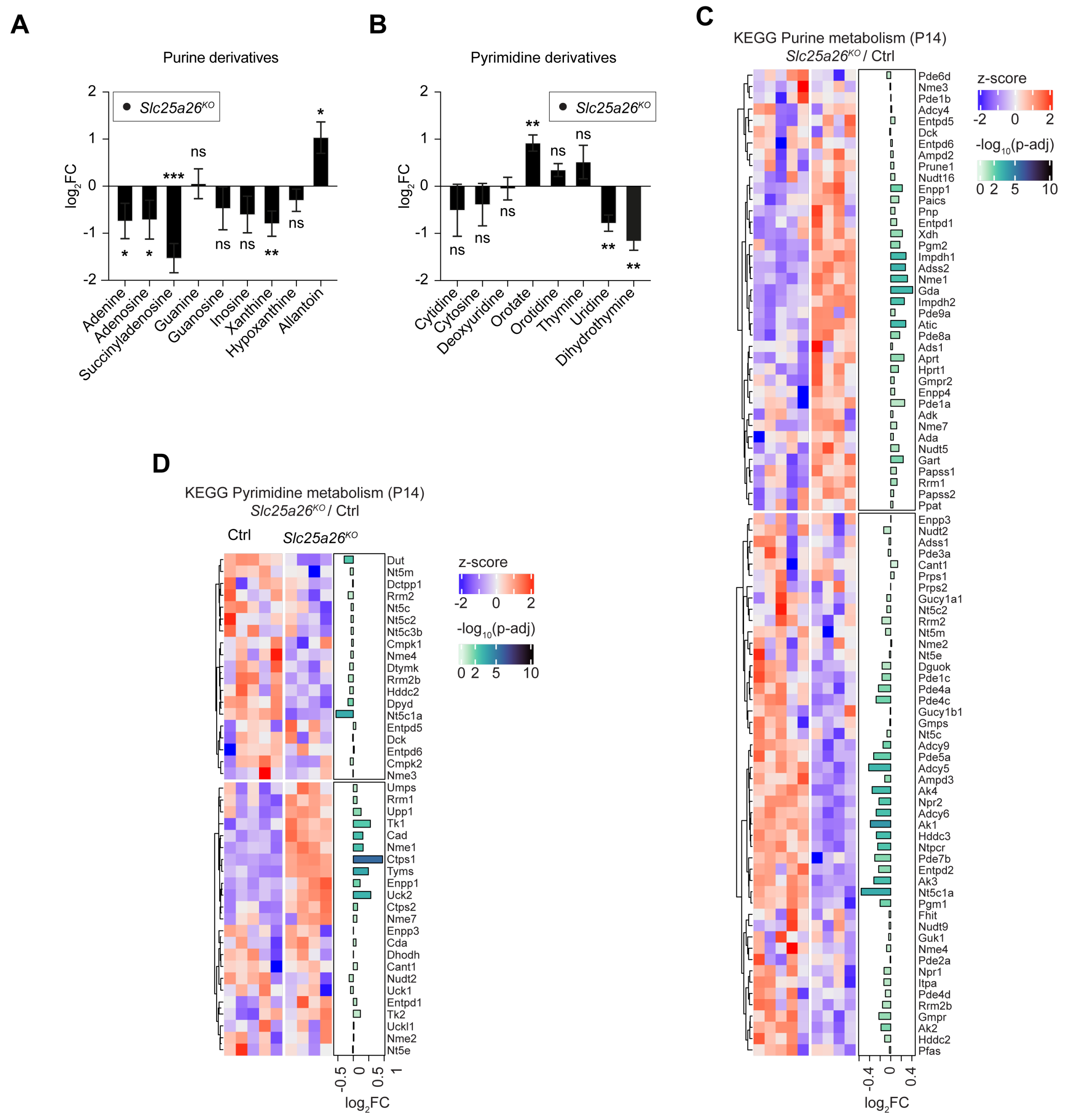


**Fig. S6. *Slc25a26* deficiency leads to nucleotide depletion, simultaneously upregulating nucleotide biosynthetic enzymes.**

(**A**) $Log_2FC$ of purine derivatives in the untargeted metabolome of *Slc25a26*$^{KO}$ hearts compared with Ctrl at P14 (n=7 for Ctrl, n=5 for *Slc25a26*$^{KO}$). (**B**) $Log_2FC$ of pyrimidine derivatives in the untargeted metabolome of *Slc25a26*$^{KO}$ hearts compared with Ctrl at P14 (n=7 for Ctrl, n=5 for *Slc25a26*$^{KO}$). (**C**) Heat map with relative z-score of heart proteomics data, showing proteins in

Purine metabolism KEGG pathway, *Slc25a26*$^{KO}$ hearts compared with Ctrl at P14 (n=5 for Ctrl, n=4 for *Slc25a26*$^{KO}$). (**D**) Heat map with relative z-score of heart proteomics data, showing proteins in Pyrimidine metabolism KEGG pathway, *Slc25a26*$^{KO}$ hearts compared with Ctrl at P14 (n=5 for Ctrl, n=4 for *Slc25a26*$^{KO}$). Data in (A and B) presented as $\log_2$FC ± s.e.m., two-tailed unpaired Student's *t*-test; ns = not significant, *$p<0.05$; **$p<0.01$; ***$p<0.001$. Significance analysis for (C and D) was performed using a linear model and moderated t-statistics. *P*-values were adjusted for multiple testing (Benjamini-Hochberg).

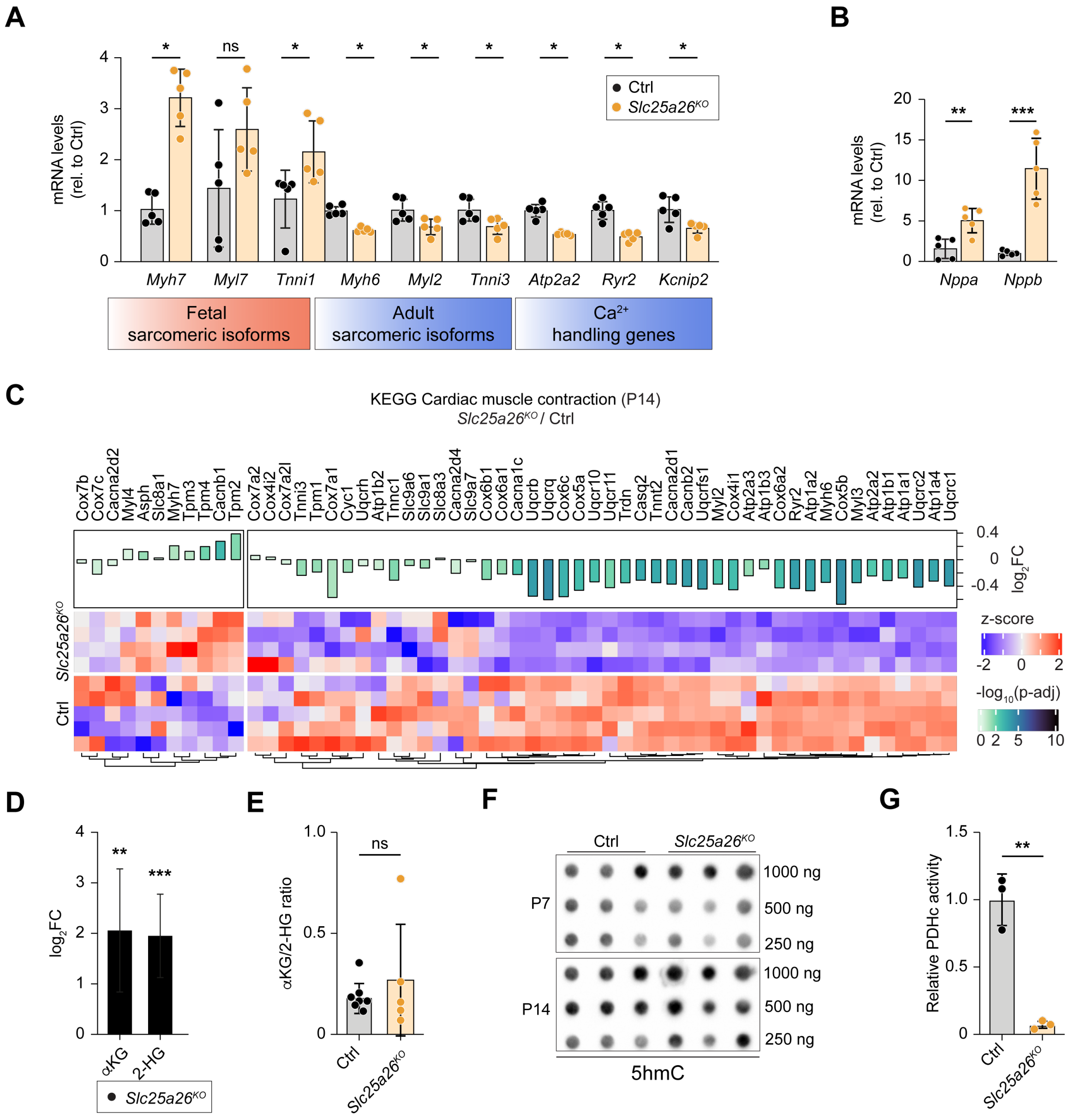


**Fig. S7. *Slc25a26* deficiency extends the cardiomyocyte proliferative window independent from global DNA demethylation.**

(**A**) Transcript levels of indicated $Ca^{2+}$ handling genes and fetal and adult sarcomeric gene isoforms in Ctrl and *Slc25a26*$^{KO}$ hearts at P14, qRT-PCR (n=5 per genotype). (**B**) Transcript levels of heart failure markers *Nppa* and *Nppb* in Ctrl and *Slc25a26*$^{KO}$ hearts at P14, qRT-PCR (n=5 per genotype). (**C**) Heat map with relative z-score of heart proteomic data, showing proteins in the Cardiac muscle contraction KEGG pathway, *Slc25a26*$^{KO}$ hearts compared with Ctrl at P14 (n=5 for Ctrl, n=4 for *Slc25a26*$^{KO}$). (**D**) $Log_2FC$ of αKG and 2-hydroxyglutarate (2-HG) in the untargeted metabolome of *Slc25a26*$^{KO}$ hearts compared with Ctrl at P14 (n=7 for Ctrl, n=5 for *Slc25a26*$^{KO}$). (**E**) Ratio of αKG and 2-HG in control and *Slc25a26*$^{KO}$ hearts at P14. (**F**) DNA dot blot analysis of 5hmC levels in Ctrl and *Slc25a26*$^{KO}$ hearts at P7 and P14 (n=3 per genotype and time point). (**G**) Pyruvate dehydrogenase activity in Ctrl and *Slc25a26*$^{KO}$ hearts at P14. (n=3 per genotype). Data in (A, B, E, G) presented as mean ± SD, each dot represents an individual mouse, two-tailed unpaired Student's *t*-test, ns = not significant, *$p<0.05$; **$p<0.01$; ***$p<0.001$. Statistical analysis for (C) was performed using a linear model and moderated *t*-statistics. *P*-values were adjusted for multiple testing (Benjamini-Hochberg). Data in (D) presented as $log_2FC$ ± s.e.m., statistical significance was determined using two-tailed unpaired Student's *t*-test; **$p<0.01$; ***$p<0.001$.

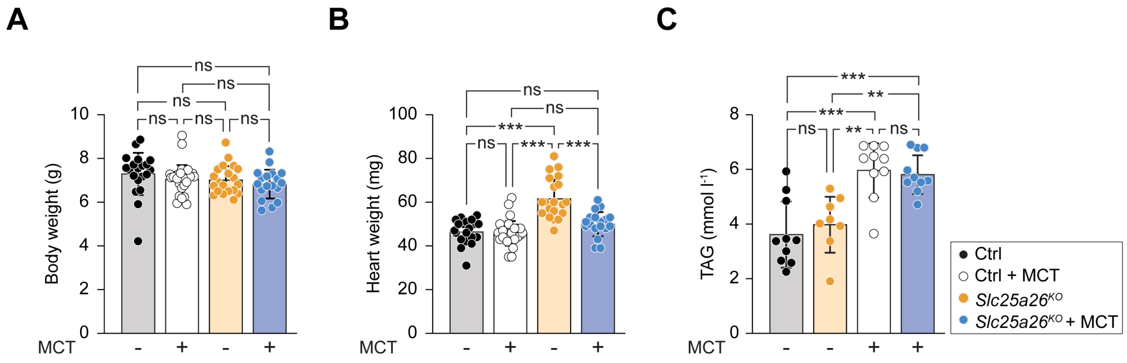


**Fig. S8. Dietary supplementation with medium-chain triglycerides (MCT) decreases cardiac mass and increases circulating blood triglycerides.**

(**A**) Body weight of Ctrl and *Slc25a26*$^{KO}$ animals fed normal or MCT-enriched chow at P14 (n=21, 26, 20, 18 for Ctrl, Ctrl+MCT, *Slc25a26*$^{KO}$, and *Slc25a26*$^{KO}$ +MCT, respectively). (**B**) Heart weight of Ctrl and *Slc25a26*$^{KO}$ animals fed normal or MCT-enriched chow at P14 (n=21, 26, 20, 18 for Ctrl, Ctrl+MCT, *Slc25a26*$^{KO}$, and *Slc25a26*$^{KO}$ +MCT, respectively). (**C**) Blood triglyceride levels in Ctrl and *Slc25a26*$^{KO}$ animals fed normal or MCT-enriched chow at P14. (n=10, 10, 8, 10 for Ctrl, Ctrl+MCT, *Slc25a26*$^{KO}$, and *Slc25a26*$^{KO}$ +MCT, respectively). Data (A–C) are presented as mean ± SD, each dot represents an individual mouse, two-way ANOVA with Tukey's multiple comparisons test; ns = not significant, **p*<0.05; ***p*<0.01; ****p*<0.001.

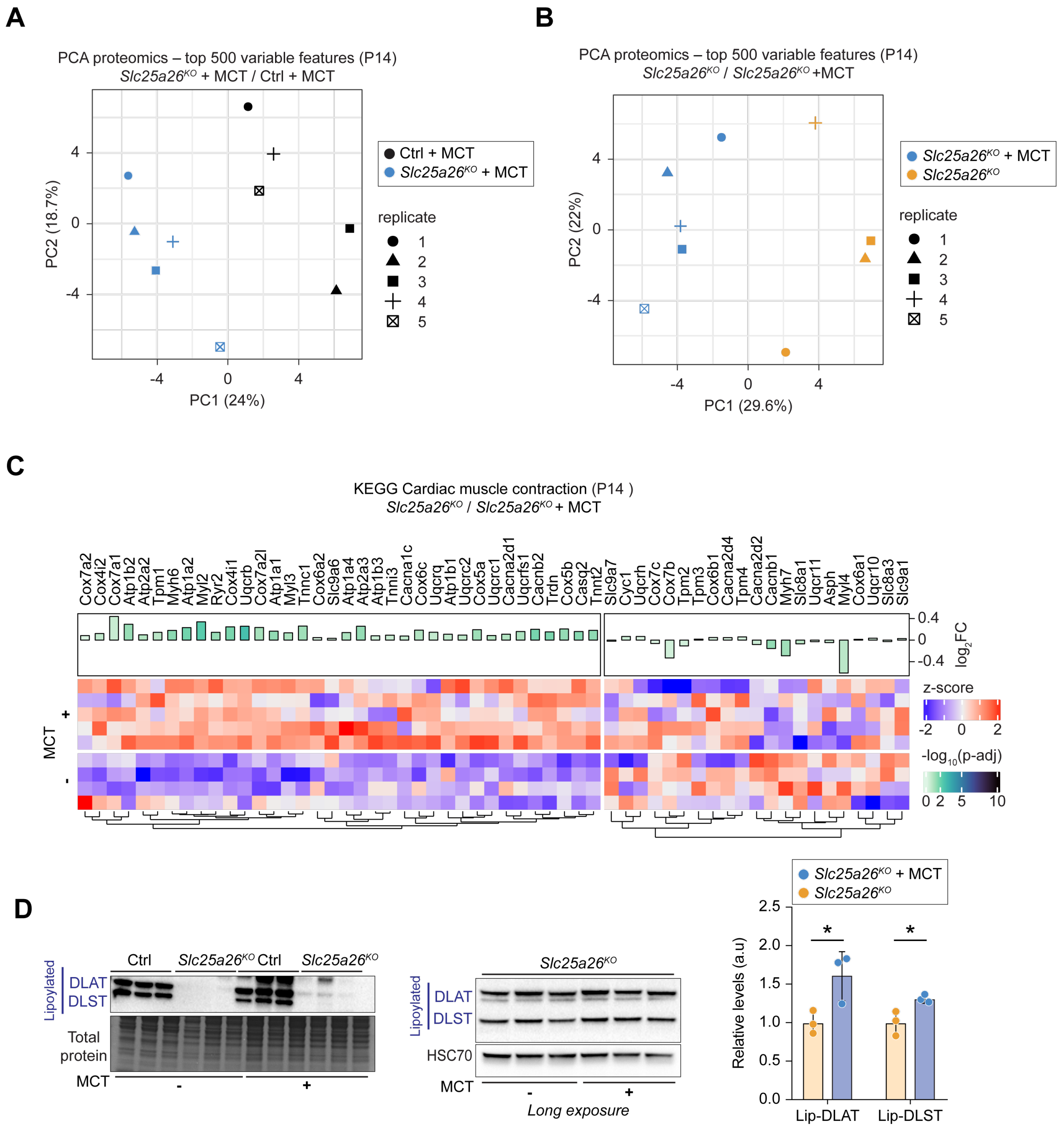


**Fig. S9. Dietary supplementation with medium-chain triglycerides (MCT) improves *Slc25a26$^{KO}$* cardiac health.**

(**A**) PCA plot of LC-MS/MS label-free whole heart proteomic data. Ctrl and *Slc25a26$^{KO}$* mice fed on MCT-enriched chow at P14 (n=5 per genotype). (**B**) PCA plot of LC-MS/MS label-free whole heart proteomic data, *Slc25a26$^{KO}$* mice fed on normal (n=4) and MCT-enriched chow (n=5) at P14. (**C**) Heat map with relative z-score of heart proteomic data, showing proteins in cardiac muscle contraction KEGG pathway from P14 *Slc25a26$^{KO}$* mice fed on normal (n=4) versus MCT-enrich (n=5) chow. (**D**) Immunoblot of P14 Ctrl and *Slc25a26$^{KO}$* heart lysates from mice fed normal or MCT-enrich chow at P14, depicting lipoylated forms of DLAT and DLST. HSC70 was used as the loading control (n=3 per genotype and treatment). Quantification of a long-exposure immunoblot of P14 *Slc25a26$^{KO}$* heart lysates fed with normal or MCT-enriched chow (n=3 per treatment) is shown on the right. Statistical analysis for (C) was performed using a linear model and moderated *t*-statistics. *P*-values were adjusted for multiple testing (Benjamini-Hochberg). Data in (D) presented as mean ± SD, each dot represents an individual mouse, two-tailed unpaired Student's *t*-test, *$p<0.05$.

**Table S1. List of TaqMan probes used in this study.**

| Probe | Source | Catalog number |
|---|---|---|
| *Slc25a26* (Mm00470957_m1) | ThermoFisher | Cat#4351372 |
| *Nppa* (Mm01255747_g1) | ThermoFisher | Cat#4331182 |
| *Nppb* Mm01255770_g1 | ThermoFisher | Cat#4331182 |
| *Myh7* Mm00600555_m1 | ThermoFisher | Cat#4331182 |
| *Myh6* Mm00440359_m1 | ThermoFisher | Cat#4331182 |
| *Myl7* (Mm00491655_m1) | ThermoFisher | Cat#4331182 |
| *Myl2* (Mm00440384_m1) | ThermoFisher | Cat#4331182 |
| *Tnni1* (Mm00502426_m1) | ThermoFisher | Cat#4331182 |
| *Tnni3* (Mm00437164_m1) | ThermoFisher | Cat#4331182 |
| *Ryr2* (Mm00465877_m1) | ThermoFisher | Cat#4331182 |
| *Atp2a2* (Mm01201431_m1) | ThermoFisher | Cat#4331182 |
| *Kcnip2* (Mm00518915_g1) | ThermoFisher | Cat#4331182 |
| *Actb* (Mm01205647_g1) | ThermoFisher | Cat#4331182 |
| *18S* (Mm03928990_g1) | ThermoFisher | Cat#4331182 |
| *Hprt* (Mm03024075_m1) | ThermoFisher | Cat#4331182 |
| *Nd1* (Mm04225274_s1) | ThermoFisher | Cat#4351370 |
| *Nd2* (Mm04225288_s1) | ThermoFisher | Cat#4351370 |
| *Nd3* (Mm04225292_g1) | ThermoFisher | Cat#4351370 |
| *Nd4L/Nd4* (Mm04225294_s1) | ThermoFisher | Cat#4351370 |
| *Nd5* (AIHSNT9) | ThermoFisher | custom made |
| *Nd6* (AIVI3E8) | ThermoFisher | custom made |
| *Cytb* (Mm04225271_g1) | ThermoFisher | Cat#4351370 |
| *Cox1* (Mm04225243_g1) | ThermoFisher | Cat#4351370 |
| *Cox2* (Mm03294838_g1) | ThermoFisher | Cat#4351370 |
| *Cox3* (Mm04225261_g1) | ThermoFisher | Cat#4351370 |
| *Atp6* (Mm03649417_g1) | ThermoFisher | Cat#4351370 |
| *Atp8* (Mm04225236_g1) | ThermoFisher | Cat#4351370 |
| *mtRnr1* (Mm04260177_s1) | ThermoFisher | Cat#4351370 |
| *mtRnr2* (Mm04260181_s1) | ThermoFisher | Cat#4351370 |
